# Light-dependent flavin redox and adduct states control the conformation and DNA binding activity of the transcription factor EL222

**DOI:** 10.1101/2024.11.06.618433

**Authors:** Aditya S. Chaudhari, Adrien Favier, Zahra Aliakbar Tehrani, Tomáš Kovaľ, Inger Andersson, Bohdan Schneider, Jan Dohnálek, Jiří Černý, Bernhard Brutscher, Gustavo Fuertes

## Abstract

The activity of the transcription factor EL222 is regulated through protein-chromophore adduct formation, interdomain dynamics, oligomerization and protein-DNA interactions, all triggered by photo-excitation of its flavin mononucleotide (FMN) cofactor. To gain molecular-level insight into the photocycle of EL222, we applied complementary methods: macromolecular X-ray crystallography (MX), nuclear magnetic resonance (NMR) spectroscopy, optical spectroscopies (infrared and UV/visible), molecular dynamics/metadynamics (MD/metaD) simulations, and protein engineering using non-canonical amino acids. The observation of only subtle atomic displacements between crystal structures of EL222 with and without blue-light back-illumination, was confirmed by NMR data indicating no major changes in secondary structure and fold compactness. Kinetic experiments in solution provided evidence for two distinct EL222 conformations (lit1 and lit2) that become sequentially populated under illumination. These two lit states were assigned to covalently-bound N_5_ protonated, and non-covalently-bound hydroquinone forms of FMN, respectively. Molecular modeling revealed differential dynamics and domain separation times arising from the three FMN states (oxidized, adduct, and reduced). Furthermore, while the dark state is largely monomeric, both lit states undergo slow monomer-dimer exchange. The photoinduced loss of α-helicity, seen by infrared difference spectroscopy, was ascribed to dimeric EL222 species. Unexpectedly, NMR revealed that all three EL222 species (dark, lit1, lit2) can associate with DNA to some extent, but only under illumination a high population of stable complexes is obtained. Overall, we propose a refined model of EL222 photo-activation where photoinduced changes in the oxidation state of FMN and thioadduct formation shift the population equilibrium towards an open conformation that favors self-association and DNA-binding.

**Significance Statement:** Flavin-binding light-oxygen-voltage (LOV) proteins constitute a prominent example of highly evolved chromophore-containing proteins that convert light into biochemical changes in the cell. However, it is not well understood how blue-light orchestrates changes in LOV structure and function. Here we show that the dynamics, oligomerization and DNA-binding properties of the photocontrolled transcription factor EL222 are dependent on both the flavin redox state and thioadduct formation. In the dark, monomeric EL222 forms transient encounter complexes with DNA. Under illumination, two distinct lit states are sequentially generated, termed lit1 and lit2, that are both able to assemble into EL222:DNA (2:1) complexes. Our results reveal the coupling between flavin photochemistry (protonation and covalent linkage) and fold stability in EL222 and potentially other flavoproteins.

## Introduction

Photoreceptors containing the light-oxygen-voltage (LOV) domain regulate multiple functions including phototropism in plants, the circadian clock in fungi, and stress response in bacteria (1-5). Bacterial LOV proteins generally feature a simple organization where the LOV sensor domain is connected to an effector/output domain via a linker sequence (3). This is the case of EL222, a transcription factor from the bacterium *Erythrobacter litoralis*, which contains an N-terminal sensory LOV domain and a C-terminal helix-turn-helix (HTH) DNA-binding domain (6). EL222 is transcriptionally inactive in the dark but avidly binds to DNA and switches on gene expression upon illumination (6, 7). Because of this property, EL222 has found widespread use in optogenetic applications that require rapid and reversible control of transcriptional activity (8-14). The LOV core domain has a classical Per-Arnt-Sim (PAS) fold, consisting of a five-stranded antiparallel β-sheet and four helical elements, one of which holds a conserved cysteine residue (Cys78 in EL222). LOV signaling is dependent on a flavin chromophore, typically flavin mononucleotide (FMN), non-covalently bound in the dark state i.e. in the absence of light stimulation. A crystal structure of dark-state EL222 shows a canonical LOV fold tightly packed against the HTH domain (“closed” conformation) and a structurally heterogeneous linker that is partially invisible, most likely disordered, and partially alpha-helical (Jα element) (6). High-resolution structure(s) of light-activated EL222 have so far been missing.

The hallmark of LOV domain photochemistry is the formation of a FMN-cysteinyl adduct, which features both a conserved cysteine residue covalently bound to the C4a atom of the FMN moiety (responsible for a shift in the absorption maximum from 450 nm to ∼390 nm), and protonation of the N_5_ atom of FMN (4, 5). Fold-switching has been proposed for EL222 with the lit state being more disordered than the dark state (15). Moreover, EL222 is mostly monomeric in the absence of illumination, while upon illumination it forms transient dimers that may be stabilized in the presence of DNA (16-18). After ceasing illumination, EL222 returns spontaneously (thermally) to the dark-state configuration in a timescale of seconds (15, 19).

In summary, according to the current model of EL222 photocycle, dark-state “closed” (LOV/HTH interactions) monomers switch to an “open” (absence of LOV/HTH interactions) monomeric adduct state that is primed for dimerization through LOV/LOV and/or HTH/HTH interactions (6, 15). Given the complex interplay between secondary, tertiary and quaternary structural changes in EL222 and the limited information gathered so far, the current model is, most likely, oversimplified.

In order to gain more insight into the structural and dynamical changes of EL222 upon illumination, both with respect to the protein chain and the FMN chromophore, we undertook an integrative approach combining macromolecular crystallography (MX), solution nuclear magnetic resonance (NMR) spectroscopy, UV/Visible and infrared (IR) spectroscopy, and molecular dynamics (MD) simulations. Importantly, all these techniques can be applied not only in the absence but also in the presence of continuous illumination, hence enabling access to photostationary conditions and minimizing sparsely populated transient species arising from pulsed illumination.

## Results

### EL222 crystal structures support adduct formation

We first determined the crystal structures of EL222 in the dark (PDB ID: 8A5R) and lit (PDB ID: 8A5S, under 470 nm illumination) states, both at 1.85 Å resolution, from crystals retrieved from the same crystallization hanging drop (**Table S1**). The overall fold closely resembles the structure determined previously (PDB ID: 3P7N (6) **Table S2**) showing the N-terminal LOV and C-terminal HTH domains connected by the long Jα helix (**Fig. 1a**). Both the dark and the lit structures are supported by continuous electron density for residues T21-A226. In addition, an N-terminal segment (residues 21-27), the C-terminal non-native residue A226 and a segment comprising residues 145-152, all absent in the previously published structure (PDB ID: 3P7N(6)) could be modeled in our electron density maps. Residues 145-149 precede the Jα helix and presumably serve as a hinge enabling relative movements of the LOV and HTH domains as part of the signaling cycle. The HTH domain fixes the position of the Jα helix in an extended conformation, hindering Jα to dock onto the LOV domain β-sheet as observed in the structures lacking the HTH domain (5). Both structures contain oxidized histidine (2-oxo-histidine) at position 101, presumably because of oxidative stress. This modification, confirmed by mass spectrometry, (**Fig. S1**) was present in EL222 prior to crystallization, likely being caused by inadvertent light or redox activation of the protein during expression and purification.

**Figure 1.**
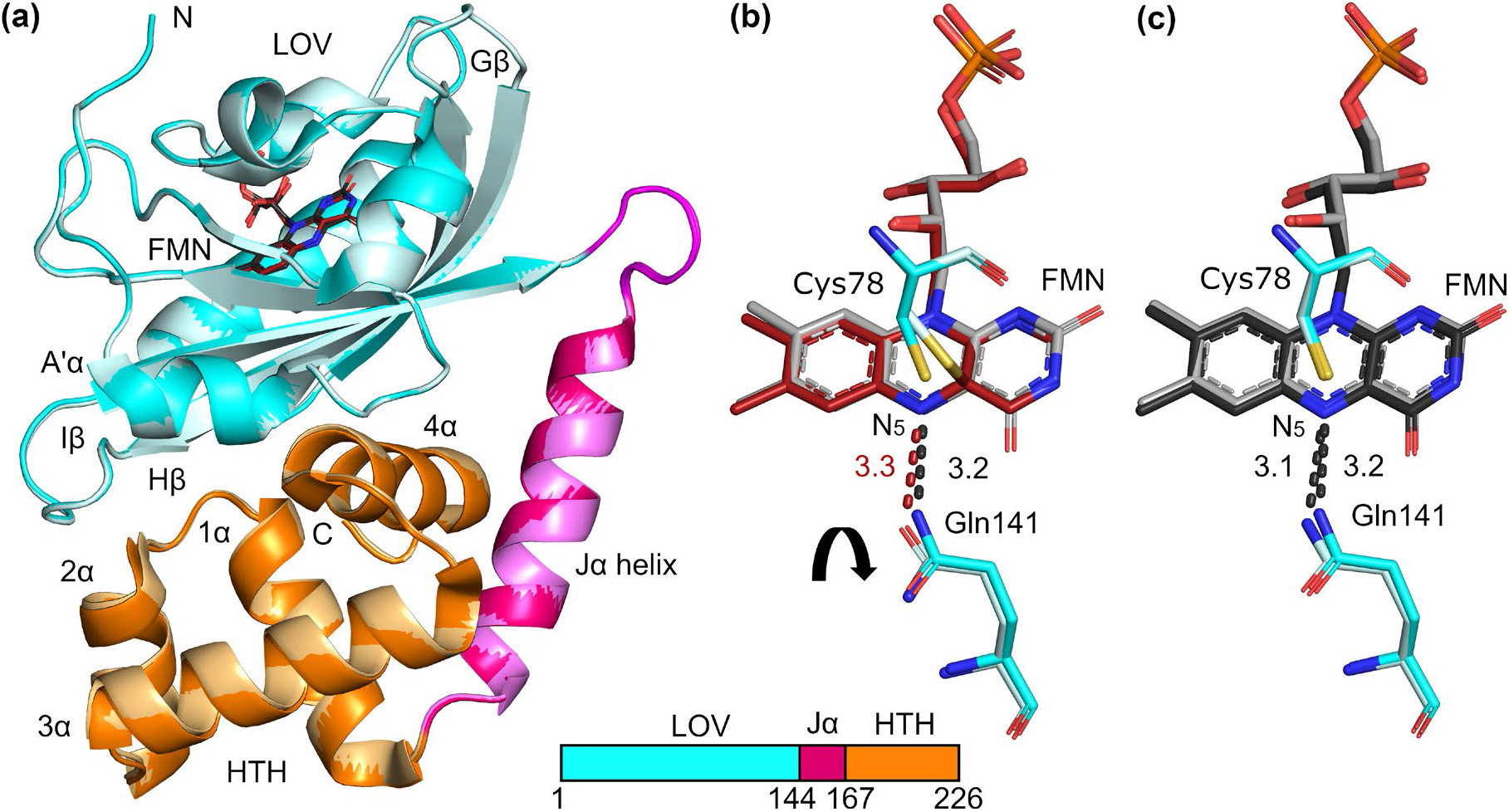
X-ray crystallography studies of EL222 show evidence for adduct formation upon illumination. Superposition of EL222 structures crystallized and measured in the dark (PDB ID: 8A5R or “dark” structure, one species) or with continuous blue light illumination (PDB ID: 8A5S or “lit” structure, containing two species). (**a**) Cartoon representation of complete EL222. The LOV domain is colored cyan, the Jα helix in magenta, and the HTH domain in orange. In the case of the “lit” structure, the color scheme is the same, but in pale shades. (**b, c**) Details of flavin mononucleotide (FMN) chromophore binding in stick representation, shown as alignment of **(b)** FMN from “dark” structure (colored gray) with FMN from “lit” structure in covalently bound state (colored red, 30% partial occupancy), and **(c)** FMN from “dark” structure (colored gray) with FMN from “lit” structure non-covalently bound (colored black, 70% partial occupancy). Residues C78 and Q141 are also shown as sticks. Hydrogen bonds of Gln141Nε2 to FMN-N_5_ in the dark and dark-like structures are shown as black dashed lines. The hydrogen bond between Gln141Oε1 and FMN-N_5_ in the adduct state is shown as a red dashed line. All distances are given in Ångströms. The rectangular inset represents the domain structure of EL222. Molecular graphics were created using PyMOL (Schrödinger, LLC).

In both states, FMN binding to the LOV domain of EL222 is similar to previous reports (5, 6) orienting the isoalloxazine C4a atom close to the Cys78S^γ^. In the dark, the C4a-S^γ^ distance refines to 3.4 Å. Under illumination, electron density around FMN indicates the presence of two populations; one is very similar to our dark state model (C4a-S^γ^ distance 3.3 Å, **Fig. 1c**). In the second population, the distance is shortened to 1.8 Å, suggesting the presence of a covalent bond (**Fig. 1b**). As a result, the C4a atom is sp3 hybridized, which in turn results in puckering of FMN around the C4a atom (**Fig. 1b**). The occupancies of the non-covalent/covalent (adduct) states of FMN in the illuminated crystals are estimated to be 70%/30%, respectively. Thus, we consider that, under our illumination conditions, 70% of the molecules in the crystal remained in the dark state and 30% were photoconverted. The arrangement of domains is very similar in the dark and lit models (**Fig. 1a**). A close inspection reveals a small, but distinguishable, shift of the HTH domain with the largest changes observed in the loop between the 1α and 2α helices, most of the 3α helix and at the beginning of helix 4α (**Fig. S2A, Table S2**). Changes in the positions of atoms in the vicinity of Gln141 (Leu138 – Ser140) propagate further along the LOV-HTH interface ultimately changing the mutual orientation of contacts between LOV and HTH domains **(Fig. S2b**).

The structural changes observed upon illumination of the crystals clearly indicate the progression of short-range atomic shifts following the formation of the FMN-Cys covalent bond. Such shifts may be the precursor to long-range conformational changes (domain separation, oligomerization). However, the latter may not occur in the whole crystal given the restrictions imposed by the crystal lattice. Thus, we next probed the light-induced structural and dynamical changes of EL222 by solution spectroscopy techniques.

### EL222 photocycle in solution is characterized by three conformational states

First, we applied solution NMR spectroscopy on uniformly ^13^C/^15^N-labelled EL222 samples. Upon illumination with blue light (488 nm), we observed significant changes in the amide ^1^H-^15^N correlation spectra (**Fig. 2a**). Notably, in addition to the dark-state species (colored black in Fig. **2a**), two lit-state populations, characterized by a distinct set of resonances, were observed and named lit1 (colored red in **Fig. 2a**) and lit2 (colored green in **Fig. 2a**). The NMR spectral signature characteristic of lit1 is detected after less than 1 min of blue-light illumination, while lit1 then slowly photoconverts into lit2 on a timescale of hours (**Fig. S3a**) under continuous illumination. The kinetic rate of the lit1-to-lit2 conversion increased linearly with the applied light power (**Fig. S3c** and **Table S3**) indicative of a single-photon photochemical process. The kinetics were also faster in the presence of ascorbate (**Fig. S3d**) indicative of a photoinduced electron transfer reaction. In the absence of light, the lit1 state reverted quickly back to the dark-state conformation, in agreement with the reported dark recovery lifetime of FMN in EL222 (∼ 40 s at pH = 6.8 and 20 °C) (15). When turning off the light, after having photoconverted EL222 to the lit2 state, samples remained largely transparent and the characteristic yellow color of oxidized FMN was not restored (*vide infra*). In addition, the NMR spectral signature did not change over time, indicating that lit2 is not spontaneously back-converting, neither to the lit1 nor to the dark state in our experimental conditions. Restoring the dark state required opening the NMR (Shigemi) tube (**Supplementary video 1**). We rationalize these observations by the requirement of molecular oxygen for the conversion from lit2 to the dark state of EL222. In the NMR tube, the water-dissolved oxygen may become depleted during the photoconversion from lit1 to lit2 states due to photoinduced electron transfer, leaving no oxidant for the back conversion to the dark state.

**Figure 2.**
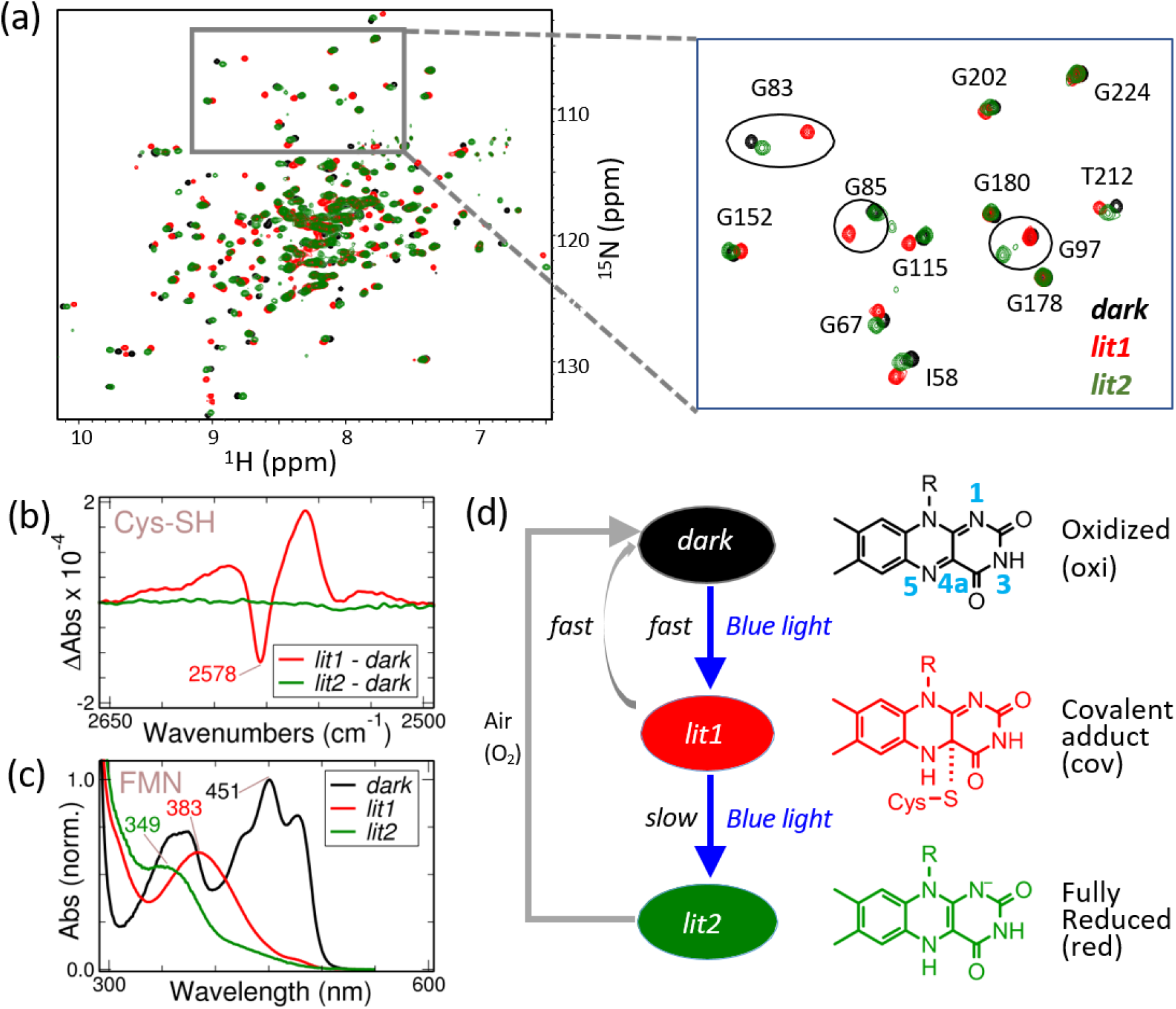
Solution spectroscopy reveals three distinct EL222 conformational states differing in the FMN redox state and adduct formation. (a) Superposition of ^1^H-^15^N correlation spectra of EL222 acquired in the dark (black), and after short (red) and prolonged (green) blue-light illumination. *Right*, small spectral region highlighting the observed chemical shift differences, indicative of changes in the electronic environment of a particular nuclear site. (b) Infrared difference spectra in the sulfhydryl stretching region of EL222. (c) UV/Visible absorbance spectra of EL222 in dark (black), lit1 (red) and lit2 (green) states. The spectrum of the lit1 state has been acquired with the variant EL222-AQTRIP, while the lit2-state spectrum has been recorded for EL222-C78A. (d) Proposed model of EL222 photocycle. In the absence of illumination, the dark state conformation with oxidized FMN prevails. Under constant illumination, two species are formed sequentially. The lit1 state (FMN-cysteinyl adduct with N_5_ protonated), which forms rapidly under blue-light irradiation, slowly (hours) converts into the lit2 state featuring fully reduced and non-covalently bound FMN. Contrary to the lit1 state, which swiftly (seconds’ time scale) reverts back to the dark state conformer upon ceasing illumination, the lit2 state is a new species that does not relax spontaneously (thermally) in the absence of oxygen.

The observed ^1^H, ^15^N chemical shift changes between the dark and the lit1 or lit2 states are mainly localized to the LOV domain (**Fig. S4**), suggesting that light-induced chemical and structural changes are sensed primarily by nuclei in the vicinity of the FMN cofactor. We therefore investigated the role of the additional HTH domain in this photocycle by repeating the NMR photoconversion experiment on the isolated LOV domain embedding the FMN cofactor. The same three photo-states (dark, lit1, and lit2) were observed for this construct (**Fig. S5a**), although with a 10-fold increase of the lit1-to-lit2 conversion rate (**Fig. S3b**). This suggests that the lit2 state may be a general property of LOV domains, while the effector (DNA binding) domain significantly slows down the conformational transition from lit1 to lit2. Previous NMR studies of a slightly different variant of EL222 assigned the ^1^H, ^13^C, and ^15^N chemical shifts in the dark state (BMRB 17640) but identified only one species upon illumination (6). By comparing the patterns in chemical shift differences (**Fig S4a** and **S4b**), we conclude that our EL222 lit1 state corresponds to the previous lit state.

To understand the nature of these two states (lit1 and lit2) we examined the redox state of the FMN moiety. Isolated FMN can undergo two successive electron reductions (eventually accompanied by proton transfer) to form either a semiquinone radical or a hydroquinone form (**Fig. S6**). While the flavin N3 is protonated in all three species, N1 only becomes protonated in the hydroquinone form at low pH, and N5 is expected to be protonated in semiquinone and hydroquinone FMN (20). Therefore, these different chemical species can be distinguished by NMR observation of their characteristic ^1^H-^15^N spectral signatures (21-23). NMR data recorded on EL222 in its different photo-states (**Fig. S7a**) are in agreement with FMN being oxidized in the dark, N5 protonated in lit1, and with all FMN nitrogens (N_1_, N_3_ and N_5_) protonated at pH = 5.5 in lit2.

To investigate whether an FMN-cysteinyl adduct is formed in the two lit states, we combined NMR, infrared and UV-Vis spectroscopy data. The ^13^C chemical shift of the FMN C4a carbon is a sensitive reporter of this adduct formation with a value of about 135 ppm for free FMN and 65 ppm if FMN is forming a covalent bond between the C4a and the sulfur atom of a nearby cysteine side-chain (22). The 2D H3(N3)C correlation spectrum, shown in **Fig. S7b**, provides an indirect measure of this NMR chemical shift with an expected H3-C4a peak doublet if no adduct is formed, and a singlet peak in case of adduct formation. These NMR data show that the chemical bond with C4a is only formed in the lit1 state. The absence of adduct formation in lit2 is further confirmed from NMR data recorded for an EL222-C78A mutant lacking the conserved cysteine that engages in covalent bond formation with the FMN. No evidence of lit1 state formation was found for this mutant, and the dark state converted rapidly to the lit2 state upon illumination (**Fig. S5b**). The same conclusion was obtained from infrared difference spectra recorded on EL222 under dark and light conditions using the SH stretching band (∼2570 cm^-1^) as a “transparent window” vibrational reporter (24, 25). The lit-minus-dark state spectrum after short irradiation, corresponding to the lit1 state, showed a negative band suggesting the disappearance of sulfhydryl bonds and their replacement by thioether bonds vibrating elsewhere (**Fig. 2b**). Interestingly, the difference spectrum after prolonged irradiation was nearly zero indicating that sulfhydryl groups are restored, i.e. the adduct is broken, in the lit2 state (**Fig. 2c**). IR difference spectra acquired for EL222-C78A in the amide I’ band showed evidence of only one lit state with minor changes in SH bond vibrations (**Fig. S8** and **S9**).

Finally, we recorded UV-visible spectra, which are well-known markers of FMN redox state and thioadduct formation (26). We also measured an EL222 mutant, named AQTrip, that has a significantly lengthened lifetime of the lit1 state in the absence of illumination (18), thus allowing us to record a UV-Vis absorption spectrum of this metastable state (**Fig. 2c** and **S10**). The absorption band around 390 nm characteristic of chemical bond formation is only visible for lit1, and the decoloring of FMN is apparent from the spectrum of lit2.

Our observations in solution can be summarized in an extended photocycle of EL222 that is depicted in **Fig. 2d**. In the absence of blue-light illumination, oxidized FMN (*oxi* FMN state), non-covalently bound to EL222, is the dominant species (corresponding to the crystal structure in the dark). Under illumination conditions, the lit1 state, which features FMN covalently bound to Cys78 and protonated at the N_5_ position (*cov* FMN state), fully reverts to the dark state in the absence of illumination. The 30% of molecules in the illuminated crystals containing an adduct most likely correspond to the lit1 state. Prolonged illumination in the absence of oxygen or illumination of EL222 variants lacking the reactive cysteine leads to the irreversible formation of lit2, which is characterized by non-covalently bound FMN in the fully (two-electron) reduced state (*red* FMN state).

### Structural and oligomerization properties of the three EL222 states

To obtain additional site-specific information about EL222 conformation and dynamics, we first performed NMR backbone ^1^H, ^15^N, ^13^CO and ^13^CA chemical shift assignments of the dark (BMRB 52621) and lit2 (BMRB 52622) states that remain stable for several days allowing the recording of a series of 3D H-N-C correlation spectra. In addition, partial (∼70%) ^1^H, ^15^N assignments were obtained for the transiently formed lit1 state (BMRB 52620). Note that our dark state NMR chemical shifts closely match those reported in (6) for a similar EL222 construct (BMRB 17640). These chemical shifts report on the local electronic environment of a nucleus, and therefore only indirectly on the molecular structure. ^1^H-^15^N chemical shift differences between the dark and lit1 or lit2 states are mainly localized in the FMN-containing LOV domain (**Fig. S4**). Backbone ^13^C chemical shifts provide sensitive reporters on secondary structural elements along the peptide sequence (27, 28). Chemical-shift based TALOS predictions (29, 30) of secondary structure propensities in EL222 are in good agreement with the structures presented here and with the previously determined crystal structure (3P7N) (6), and no significant changes in secondary structure are observed upon prolonged blue light illumination (lit2) (**Fig. S11**). Furthermore, ^15^N transverse relaxation times (T_2_) measured for individual backbone amides along the peptide sequence in the three photo-stationary states indicate that monomeric EL222 forms a compact protein structure in all states (**Fig. 3a**). No indication of additional local dynamics or segmental motion of the individual protein domains upon illumination is obtained from these data. The rotational tumbling correlation times extracted from the measured ^15^N T_2_ and additional TRACT data (**Fig. S12**) are very similar for the dark and lit states, and in agreement with monomeric EL222 particles in solution. However, these results do not exclude the possibility of EL222 dynamics occurring on longer time scales.

**Figure 3.**
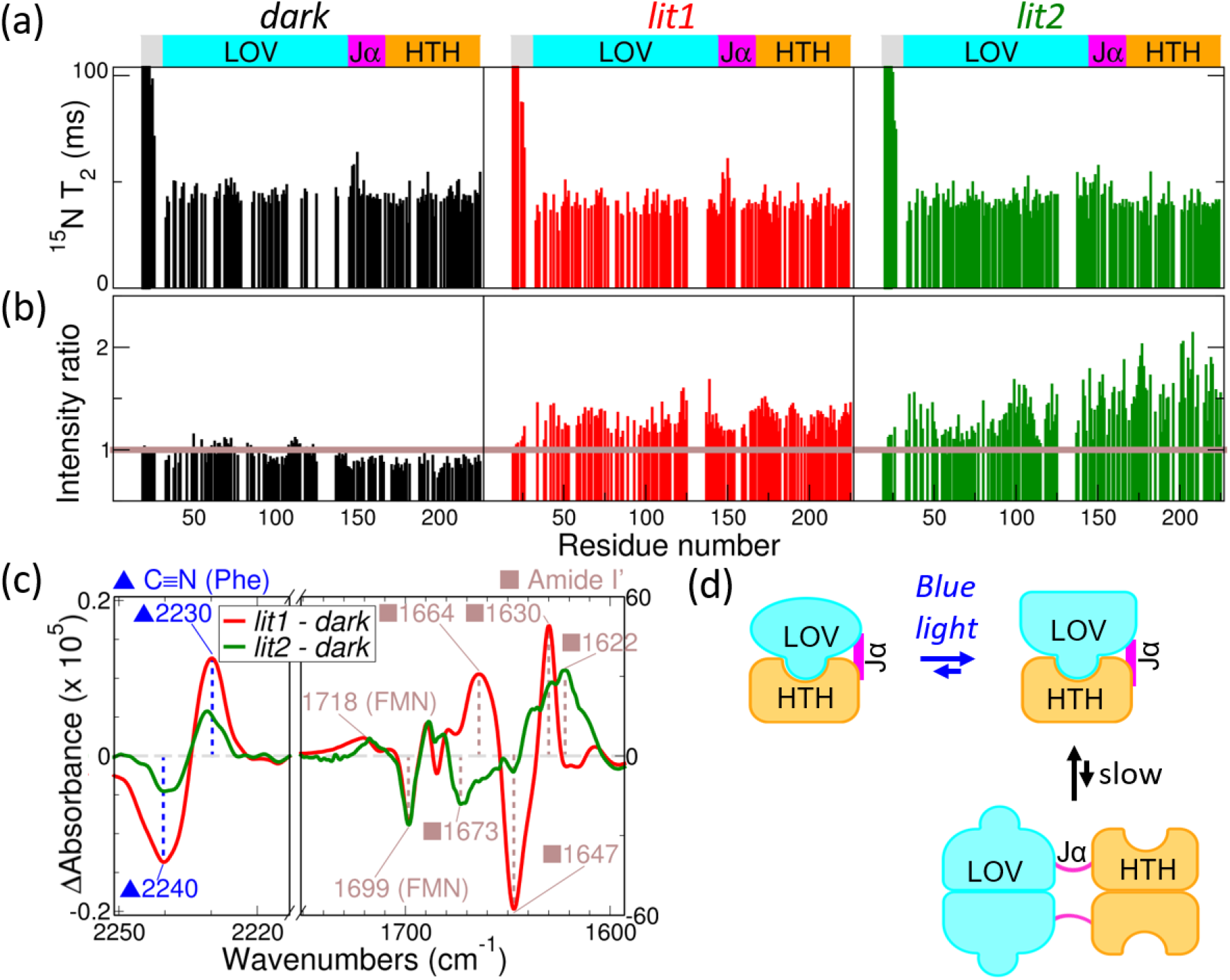
The three conformational states of EL222 differ in oligomerization propensity. (a) Residue-specific transverse ^15^N backbone relaxation times (T_2_) plotted as a function of the EL222 sequence for the dark (left), lit1 (middle), and lit2 (right) states. (b) NMR intensity ratios of the different states plotted as a function of residue number upon 4-fold EL222 dilution from 120 μM to 30 μM. (c) Infrared difference spectra in the nitrile (C≡N) stretching region of EL222-W31CNF (*left*), and in the amide I’ (C=O) region of EL222-WT (*right*) support the presence of two conformational states under continuous illumination differing in oligomerization and secondary structure content. The more pronounced and clear changes affect the lit1 state, which can be ascribed to loss of α-helicity and gain of disorder with respect to the dark state. In the case of lit2, the secondary structure changes may involve restructuring of β-sheets. (d) Model of photoinduced conformational changes of EL222 in the absence of DNA: Blue-light promotes EL222 oligomerization, with the fraction of molecules that remain monomeric showing virtually no secondary structural changes with respect to the dark state. Only some residues in EL222 dimers (NMR-invisible) undergo helix-to-coil transition, most likely in the connecting Jα element.

IR difference spectra in the secondary structure-sensitive amide I’ band (1600-1700 cm^-1^), mainly reporting on the C=O stretching vibrations of the protein backbone (31), suggest that the two light-induced conformational states of EL222 differ in their secondary structure content (**Fig. 3c** and **S9**, and **Table S3**). While the lit1 state shows a loss of α-helical content with respect to the dark state (negative band at 1647 cm^-1^), as previously reported (15), no such drop of α-helicity is detected for the lit2 state. Moreover, the negative/positive bands at 1673 cm^-1^/1622 cm^-1^, which may be tentatively assigned to β-sheets, suggest that the secondary structure content of lit2 is different from the dark state. This apparent discrepancy between NMR (no major secondary structure changes) and IR data (distinct secondary structure in dark, lit1 and lit2 states) can be reconciled considering a model where monomeric EL222 is in slow exchange (rate constant < 100 s^-1^) with oligomeric species. While EL222 dimers, and higher-order oligomers, are essentially NMR-invisible under our experimental conditions, IR is sensitive to all species present in solution regardless of their size. Thus, photoinduced secondary structure changes seem to occur preferentially in oligomeric EL222 species.

To further investigate potential differences in oligomerization propensity between the dark and lit states of EL222, we have performed additional NMR and IR experiments. As oligomerization is a concentration-dependent process, we expect that more oligomers are formed at higher protein concentration. With the hypothesis of a slow exchange process, we expect the intensity of NMR signals arising from monomeric EL222 species to be reduced at higher EL222 concentrations. The peak intensity ratios measured in ^1^H-^15^N spectra after 4-fold protein dilution are plotted in **Fig. 3b**. While in the dark no significant change in peak intensities is observed, the relative peak intensity increases upon dilution for both the lit1 and lit2 states, indicative of the presence of oligomeric species. For IR spectroscopy, we employed an EL222 variant carrying the non-canonical amino acid 4-cyanophenylalanine (CNF) at position 31. This variant was previously shown to report on the monomer-multimer equilibrium via a red-shift (the nitrile band of oligomeric species vibrating at lower frequency than the monomers) (15). The IR difference spectra of EL222-W31CNF in the C≡N stretching region are similar for lit1 and lit2 states (**Fig. 3c** and **S9**, and **Table S3**) again suggesting the presence of oligomers in both cases. The smaller spectral amplitudes of the lit2 state may be compatible with a lower population of oligomers relative to lit1. These observations support a model (**Fig. 3d**) where, in the presence of blue light, monomeric EL222 is in slow exchange with oligomeric species, most likely due to interaction of EL222 molecules in an “open” conformation, where the LOV domain is no longer protected by the HTH domain from binding to another LOV domain. This model is consistent with photoinduced tertiary structure changes in EL222 observed earlier by hydrogen-deuterium exchange and proteolysis susceptibility experiments (6).

### Molecular modeling indicates FMN-dependent dynamics of EL222

To gain deeper insights into the structural dynamics of EL222, a series of MD simulations were conducted (see details in **SI**). All-atom molecular models of EL222 were created featuring FMN in its *oxi, cov*, and *red* states (**Fig. 4a**), intended to reproduce the properties of the experimentally determined dark, lit1, and lit2 states, respectively. Specifically, the *oxi* model contains oxidized FMN non-covalently bound to EL222; the *cov* model features FMN covalently bound to the Cys78 residue, with a hydrogen at position N5 (N_5_H), and a 180° rotated Gln141 residue; and the *red* model represents fully reduced, negatively charged FMN (N_5_H/N_1-_) non-covalently-bound to EL222 (**Fig. 4a** and **Table S4**).

**Figure 4.**
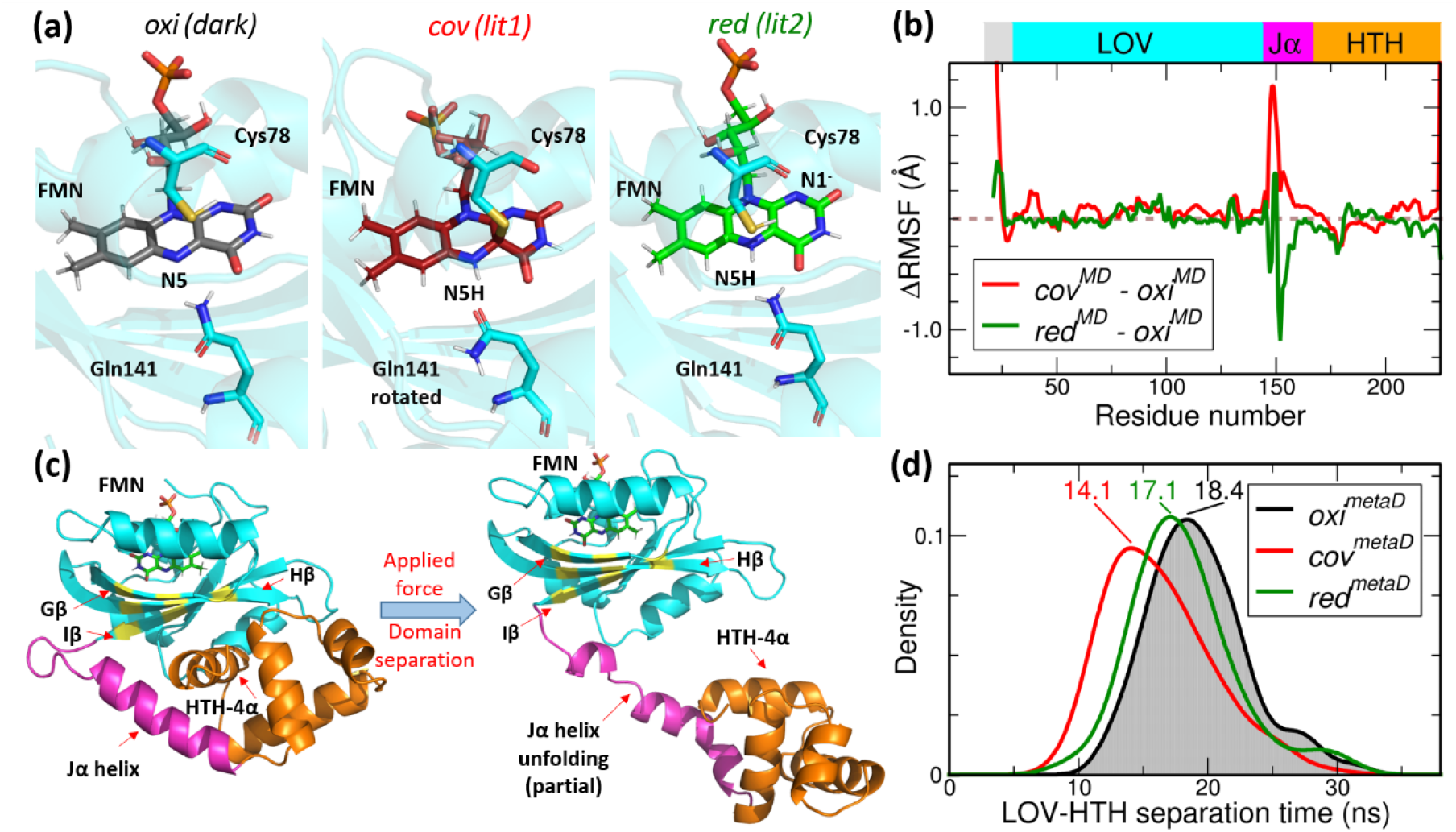
Molecular modelings suggest differential dynamics and “opening” propensities of the three EL222 states. (a) Molecular models of oxidized (*oxi*) FMN (dark), covalently-bound (*cov*) FMN (lit1), and fully reduced (*red*) FMN (lit2) used for molecular dynamics (MD) simulations, highlighting chemical differences in the FMN binding pocket of EL222 due to (de)protonation, adduct formation, and glutamine rotation. Relevant amino acids in the LOV binding site (Cys78 and Gln141) are depicted as cyan sticks. (b) Differential root mean square fluctuation (ΔRMSF) plot of Cα atoms from 3 μs cumulative molecular dynamics (MD) simulations of *cov*^*MD*^ and *red*^*MD*^ relative to *oxi*^*MD*^. (c) Representation of LOV-HTH domain separation in EL222 employing metadynamics (metaD) with a bias potential applied along an inter-domain distance defined as the distance between the center-of-mass of HTH-4α residues (shown as an orange helix) and the center-of-mass of selected residues in Iβ, Hβ, and Gβ regions (shown as yellow beta strands). The linker (including the Iβ-Jα loop and the Jα-helix) and FMN chromophore are shown in magenta and green, respectively. (d) Density plots of LOV-HTH separation time distributions for separation extracted from metaD simulations (160 replicas * 50 ns length). Mode values for domain separation time (in nanoseconds) of *oxi*^*metaD*^, *cov*^*meta*D^ and *red*^*metaD*^ are indicted for each distribution.

Employing Root Mean Square Fluctuation (RMSF) analysis, which evaluates the average displacement of atomic positions relative to their mean positions throughout the MD simulation, we systematically investigated flexible and rigid structural regions, including α-helices and β-strands, within the EL222 protein. The differential RMSF (ΔRMSF) profiles of all Cα atoms of EL222 protein residues in *cov*^*MD*^ and *red*^*MD*^ simulations, relative to *oxi*^*MD*^, are depicted in **Fig. 4b** (individual RMSF plots can be found in **Fig. S13**). Most regions of EL222 have low ΔRMSF values, indicating that the protein remains quite stable, regardless of the FMN state. The N-terminal residues exhibit the most pronounced variations of RMSF in *cov*^*MD*^ and *red*^*MD*^ trajectories compared to *oxi*^*MD*^. Enhanced RMSF and flexibility relative to *oxi*^*MD*^ are observed in the Iβ-Jα loop and the N-terminal part of Jα-helix (residues Asp147-Arg155) of the *cov*^*MD*^ simulations. This might indicate a conformational change or increased flexibility due to covalent bond formation in the LOV domain.

Although these results offer valuable insights into the dynamics of EL222, our classical MD simulations do not provide information on the key photoinduced processes, such as the detachment/unfolding of the Jα-helix region, or domain rearrangements (32). To explore the dissociation of LOV and HTH domains in EL222, we employed metadynamics (metaD) simulations (33-35) of the three molecular models (*oxi, cov*, and *red*, **Fig. 4a**) by applying a biased potential along an inter-domain distance that consequently induces partial disordering of Jα (**Fig. 4c**, see **SI** for more technical information). The time when this applied force effectively caused inter-domain separation was collected for *oxi*^*metaD*^, *cov*^*metaD*^, and *red*^*metaD*^ (**Fig. S14a**). The density plots of measured LOV-HTH domain separation times for EL222 reveal significant differences among the various FMN states (**Fig. 4d**, the corresponding histograms are presented in **Fig. S14b**). The most common (mode) separation time in the simulation of the oxidized state (*oxi*^*metaD*^) is 18.4 ns, displaying a relatively normal distribution. Conversely, the simulation of the covalent state (*cov*^*metaD*^) shows a shorter separation time of 14.1 ns, indicating that adduct formation and protonation at FMN-N5 accelerate domain separation. The simulation of the reduced N1^-^ state (*red*^*metaD*^) reports a mode time of 17.1 ns, intermediate between *oxi*^*metaD*^ and *cov*^metaD^. The marked decrease of LOV-HTH separation times in *cov*^*metaD*^ suggests that the *cov* configuration enhances structural flexibility (**Fig. 4d**).

In conclusion, the observed increased fluctuations and shorter domain dissociation time in *cov* models suggests that Cys78-FMN adduct formation and N5 protonation enable large scale conformational changes of EL222. *Red* forms displayed mixed features of *ox*i and *cov* states, showing reduced fluctuations and intermediate domain separation dynamics.

### DNA binding of EL222 in the dark and lit states

Differences in DNA binding between dark and light conditions are supposed to be at the center of transcriptional regulation by EL222 in the cell. Therefore, as a next step, we investigated *in vitro* DNA binding of EL222 in the dark, lit1, and lit2 states by NMR spectroscopy. We chose a nanomolar-affinity double-stranded oligonucleotide of 33 base pairs (DNA33) containing the 12-bp recognition motif previously identified by SELEX methods (7).

*DNA binding of EL222 in the dark*. When adding increasing amounts of DNA33 to a ^13^C/^15^N-labeled EL222 sample in the dark, we observed changes in the ^1^H-^15^N correlation spectra revealing an interaction between DNA33 and EL222 even in the absence of light. The first observation is that a number of amide sites show linear peak shifts with increasing amounts of DNA added (**Fig. 5a**). This indicates a fast exchange process between free and DNA-bound EL222. In the following, we will refer to this fast exchanging EL222-DNA species as the encounter complex. Similar peak shifts are also observed when using different overall concentrations in equimolar EL222:DNA mixtures (**Fig. 5b**), as expected for an inter-molecular association. The DNA-binding site of EL222 in this encounter complex can be inferred from the computed ^1^H/^15^N chemical shift changes upon DNA binding (**Fig. S15a**). Displaying the largest shifts on the crystal structure of EL222 (**Fig. S15b**) revealed that a large part of the HTH domain as well as a helical segment of the LOV domain (Fα) are involved in this transient interaction with DNA. These DNA-induced chemical shift changes correlate with positive surface patches of EL222 (**Fig. S15c**) suggesting that EL222-DNA encounter complex formation is electrostatically-driven. The second observation from these NMR titration data was a loss in peak intensity upon DNA binding for almost all amide sites in EL222 (**Fig. S15d**). An exception is the highly flexible N-terminal end (Asp19-Asp20-Thr21-Arg22), for which the intensity decreases gradually towards the structured part of the protein, with Asp19 being almost unaffected (**Fig. S16a**). These data can be rationalized by the formation of an EL222-DNA complex that is invisible in the ^1^H-^15^N spectra, except for the N-terminal residues that remain highly flexible and thus observable by NMR (**fig. S16b**). The observed intensity loss is not uniform across the structured part of EL222. The signal loss is particularly pronounced in spectral regions affected by encounter complex formation (HTH domain) where, in addition to signal loss due to complex formation, these amide sites are also affected by conformational exchange line broadening due to chemical shift modulation on the micro-to millisecond time scale. The average intensity decrease in the LOV domain provides an estimate of the percentage of EL222 molecules that are involved in complex formation, e.g. ∼30% for the 2:1 mixture (indicated by bars in **Fig. S15d**).

**Figure 5.**
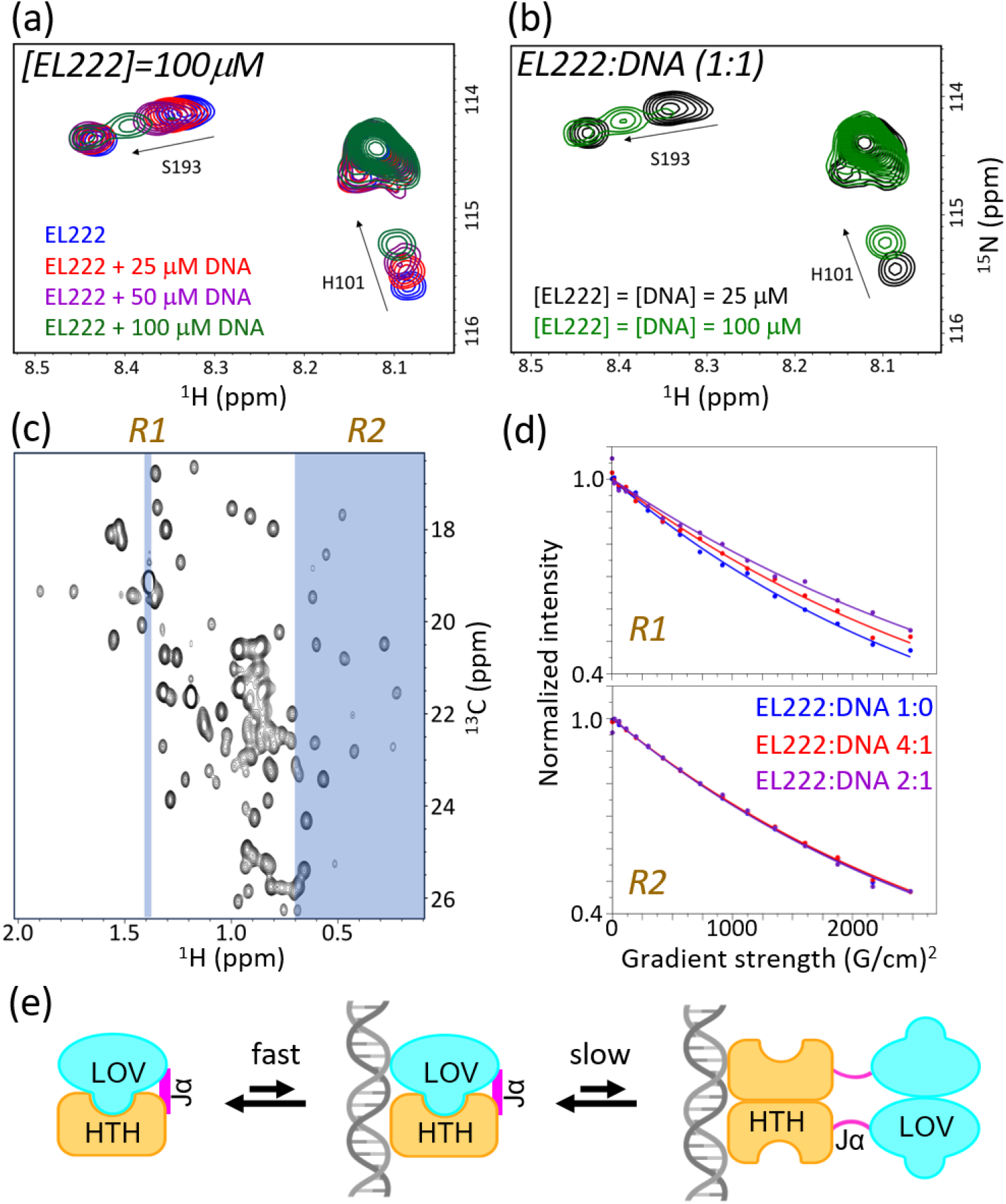
EL222 interacts with DNA in the absence of light. (a) ^1^H-^15^N NMR spectral region of EL222: DNA mixtures highlighting two residues that show linear chemical shift changes with increasing DNA concentration. (b) The same ^1^H-^15^N NMR spectral region highlighting the effect of total concentration of equimolar EL222-C78A mixtures on the NMR chemical shifts. (c) Methyl region of ^1^H-^13^C spectra of EL222. The two ^1^H spectral regions highlighted in blue correspond to a highly flexible methyl group in the monomer and complex (R1), and a number of well-structured methyl groups located in the hydrophobic protein core (R2). (d) 1D methyl 1H DOSY NMR data. The NMR intensity, integrated over the spectral region (R1 or R2) is plotted as a function of gradient strength. (e) Model of EL222-DNA interactions in the dark. Monomeric EL222 is in fast exchange between DNA-free and DNA-bound (monomeric encounter complex) forms. This encounter complex is in slow exchange with EL222-DNA complexes of 2:1 stoichiometry.

The fact that NMR signals of highly flexible sites in the EL222-DNA complex, e.g. N-terminal residues, remain NMR-observable can be exploited to obtain quantitative information on the complex stoichiometry by NMR translational diffusion measurements. We have thus performed ^1^H amide and ^1^H methyl DOSY experiments for different EL222:DNA mixtures. While both, amide and methyl-based experiments resulted in similar results, the methyl spectra were of better quality and thus used for further quantification. Two distinct methyl ^1^H spectral regions (R1 and R2) have been defined for the quantification of translational diffusion properties (**Fig. 5c**): R1 reports on the population weighted average diffusion properties of free EL222 and the EL222-DNA complex (highly flexible methyl group), while R2 is only sensing the diffusion of monomeric EL222. As expected, the diffusion curves extracted for the R2 region are independent of DNA concentration, while for the R1 region diffusion slows down with increasing amounts of added DNA (**Fig. 5d**). A quantitative analysis of these data is in agreement with a stoichiometry of two EL222 molecules binding one DNA33 molecule, as expected from literature data (6, 7, 16). Our NMR data on DNA binding by EL222 in the dark can be summarized by the kinetic scheme shown in **Fig. 5e**.

*DNA binding of EL222 in the lit states*. Finally, we investigated the effect of light on EL222-DNA complex formation by NMR and IR spectroscopies. In order to be able to distinguish the DNA binding capabilities of the two lit states, we recorded ^1^H-^15^N correlation spectra of EL222-WT (**Fig. 6a**) and EL222-C78A (**Fig. 6b**) in a 2:1 EL222:DNA mixture under continuous low-power blue-light illumination. Under these conditions, the EL222-WT spectra report on lit1 state properties, while the EL222 mutant lacking Cys78 photo-switches directly to lit2, thus providing a convenient way of probing DNA binding in the lit2 state. In both cases, the NMR signal intensity (except for the N-terminal tail) is further reduced, indicative of a shift in the population equilibrium towards the EL222-DNA complex. Interestingly, 100% complex formation is only observed for the lit1 state. In the lit2 state, about 25% of the EL222 molecules remain in an exchange between monomeric EL222 and the encounter complex. The IR double difference spectra of EL222:DNA 2:1 mixtures suggest DNA-dependent conformational changes in secondary structure (amide I’ band reporter) and oligomerization (CNF reporter), which probably arise from the NMR-invisible species (**Fig. 6c**). In other words, free and DNA-bound EL222 dimers are structurally different. Although the lit1 and lit2 spectra are qualitatively similar, the observed amplitude and frequency shifts indicate structural differences between the two lit states. Overall, we can extract two novel conclusions: 1) covalent bond formation is not essential for DNA binding but its presence increases the probability of the interaction between EL222 and DNA, 2) FMN reduction is also not essential for DNA binding but necessary to shift the equilibrium towards long-lived EL222:DNA (2:1) complexes (**Fig. 6d**). Therefore, EL222 domain separation, dimerization, and DNA binding are coupled processes that can be influenced by light, which changes the free energy of individual states, thus shifting the population equilibria as illustrated in **Fig. 7**.

**Figure 6.**
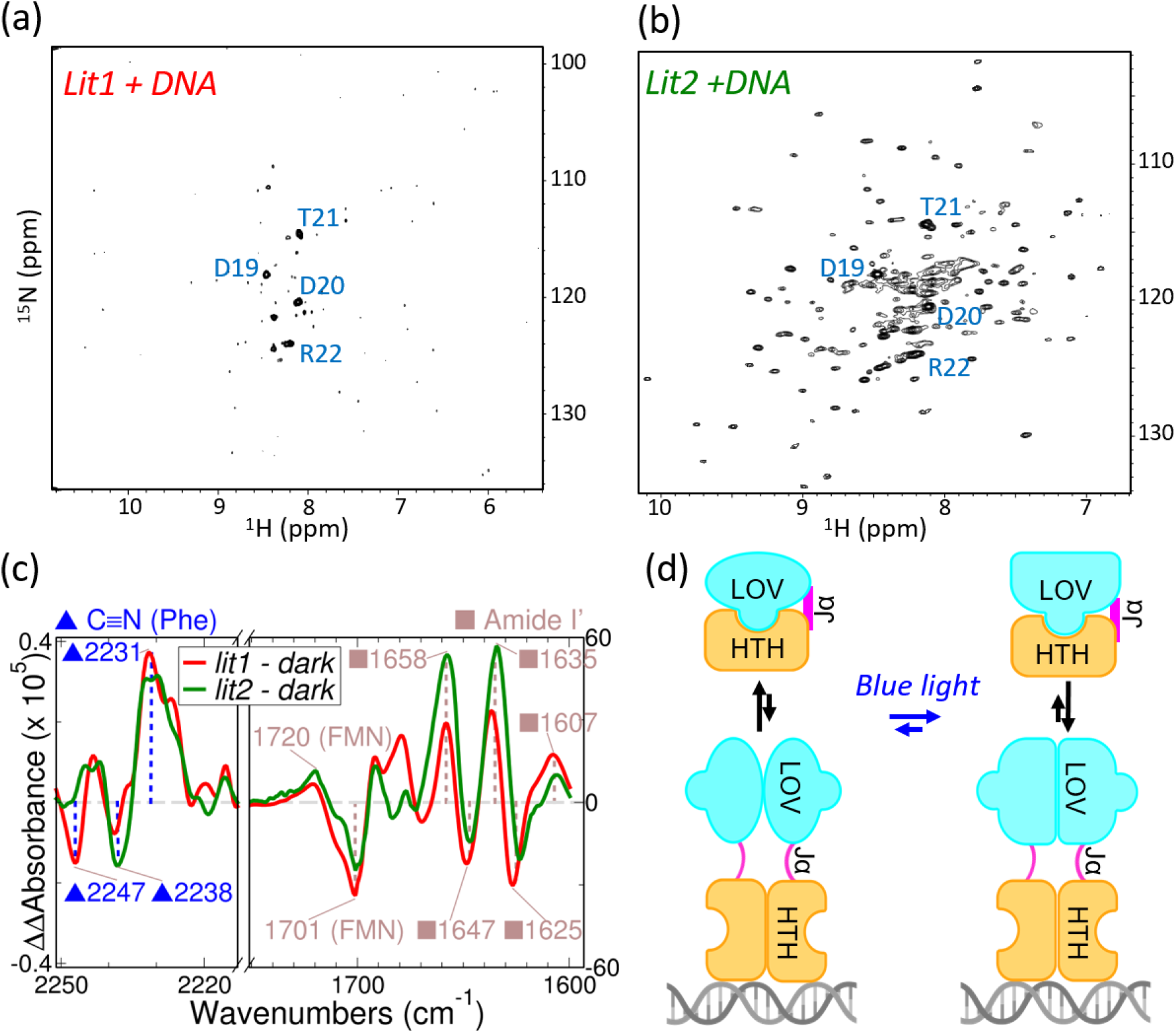
The two lit states interact with DNA with distinct efficiencies and lifetimes. ^1^H-^15^N correlation spectra of 2:1 EL222:DNA mixtures of (a) EL222-WT (100 μM) and (b) EL222-C78A (100 μM) recorded under weak blue light illumination. The flexible N-terminal residues that remain visible upon complex formation are annotated. (c) Double difference infrared spectra of EL222-W31CNF in the nitrile (C≡N) stretching region (*left*) and EL222-WT in the amide I’ (C=O) region (*right*) calculated as the difference between the light-minus-dark spectra in the presence minus the absence of DNA. (d) Model of EL222-DNA interactions upon blue-light illumination. Light shifts the binding equilibrium towards DNA-bound EL222 dimers. Elliptical and rectangular shapes denote the slightly different conformation of the LOV domain arising from changes in the redox/protonation state of FMN and presence/absence of a covalent bond.

**Figure 7.**
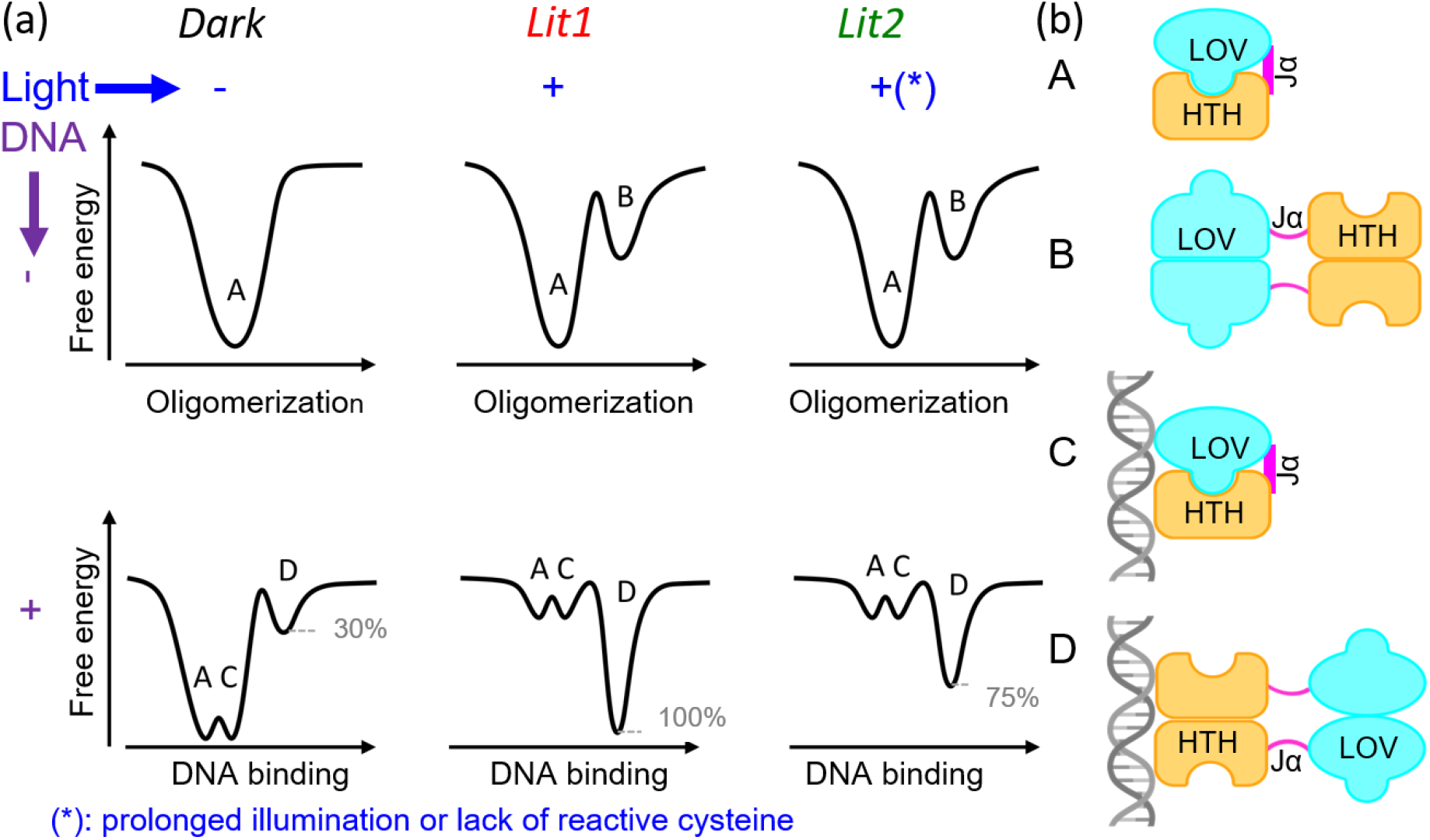
EL222 free-energy landscape model of photoinduced oligomerization and DNA binding. (a) In the absence of DNA, the dark state is characterized by a nearly 100% monomeric conformation (A), while the lit1 and lit2 states feature a mixture of monomeric (A) and dimeric (B) species. In the dark, EL222 transiently binds to DNA as monomer (C species), and also forms a small amount of 2:1 EL222:DNA complex (D species). In presence of blue-light, EL222 oligomerization and DNA binding is significantly enhanced with the largest population shift observed for lit1. (b) Proposed structural models of A (free monomer), B (free oligomer), C (DNA-bound monomer), and D (DNA-bound oligomer) species. Note that the exact conformation of A, B, C and D species will be slightly different in the dark (oxidized FMN non-covalently bound), lit1 (FMN protonated at N_5_ and covalently bound), and lit2 (fully reduced and non-covalently bound FMN) states.

## Discussion

By employing a combination of complementary methods in solution (NMR, IR, UV/vis spectroscopy), *in silico* (MD and metaD simulations), and *in crystallo* (MX), our study provides novel insights into the structural and dynamical changes that EL222 undergoes during its photocycle. The different redox/adduct states of FMN (*oxi, cov*, and *red*) give rise to three EL222 states (dark, lit1, and lit2) that show distinct behaviors with respect to their conformational dynamics (domain opening), oligomerization propensity, and DNA binding ability.

Our lit1 state corresponds to the previously described states of LOV proteins obtained after continuous illumination for short times, referred to as lit (6, 36), light (37), signaling (38), A390(39) or adduct (40) state. The distinctive feature of lit1 is the presence of a covalent bond between FMN and the active-site cysteine which is breakable in the dark. In addition, we could structurally and dynamically characterize a new photo-induced state of EL222 bearing fully reduced FMN, named lit2 which, to the best of our knowledge, has not been described before. However, LOV domains containing fully reduced FMN have been previously detected by UV/Visible spectroscopy after prolonged photoexcitation (37) or upon chemical reduction with sodium dithionite (38). Lit2 has mixed features of dark and lit1 states. On the one hand, dark and lit2 states have no covalent bond between FMN and Cys78. On the other hand, both lit1 and lit2 share an enhanced oligomerization propensity and the ability to form 2:1 complexes with DNA. One significant difference between lit1 and lit2 is that only lit1 can form 100% EL222:DNA complexes after mixing 2 moles of EL222 with 1 mole of DNA. Adduct scission in the lit2 state is compatible with the metastability and reversibility of the C-S bond (41).

The solution lit2 state of EL222 may have been previously undetected due to different irradiation conditions. Here we were applying continuous illumination in an airtight Shigemi tube, while pulsed illumination in a standard 5 mm NMR tube was used in previous work (6). The combination of prolonged illumination and minimal oxygen exchange are key for the photoconversion from lit1 to lit2 states. Because of the requirement of high irradiances and anaerobic conditions, we do not believe that the lit2 state can be readily formed from “canonical” LOV photosensors, i.e. those containing the reactive cysteine, under common environmental conditions. However, LOV photoreceptors carrying alternative residues at that position, e.g. histidine, have been described (42). Thus, the lit2 state may be the prevalent species in these “non-canonical” LOV proteins upon blue light absorption.

We cannot exclude that the lit2 state is also impacted by oxidative damage due to prolonged light exposure (43, 44), potentially leading to protein aggregation (45). Indeed, flavin-binding fluorescent proteins derived from the LOV domain have been engineered as genetically encoded photosensitizers that generate toxic reactive oxygen species (ROS) (46). We have evidence for the oxidative modification of certain amino acids in EL222, in particular the 2-oxo-histidine residue found in the crystal structures. Such a modification was previously observed in structurally similar proteins such as miniSOG (PDB ID: 6GPV), a flavin-containing protein developed from the LOV2 domain of *Arabidopsis thaliana* phototropin 2 specifically to produce singlet oxygen upon illumination (47). The increase in ROS production has been associated to the release of FMN to the solvent (43), which we do not observe in our experiments. Thus, although oxidized residues may be found in the lit2 state, it is unlikely that they become the dominant factor responsible for the conformational changes. Instead, we argue that the main driving force underlying the formation of the lit2 state is the loss of the FMN-Cys covalent bonding and full reduction of the embedded FMN chromophore. Massive LOV aggregation, i.e. clustering, has only been reported for two LOV proteins under certain conditions (45, 48). Most LOV domains undergo reversible oligomerization up to tetramers. For instance, RsLOV undergoes light-induced dimer-to-monomer transitions (49), while in AtLOV1 dark-state dimers assemble into lit-state tetramers (50). Similar to EL222, light-induced dimerization is also part of the activation mechanism of VIVID (51) and aureochrome-1 (52). In line with the latter studies, we also found that EL222 undergoes reversible oligomerization from both lit1 and lit2 states, although the exact stoichiometry remains unknown.

The slow kinetics of lit1-to-lit2 photoconversion in EL222-WT (time scale of hours) contrasts with the fast formation of lit2 in EL222-C78A (less than one minute), and with the ultrafast primary flavin photoreactions (sub-second time scale). Sub-millisecond LOV domain photochemistry is essentially the same for all LOV variants with only minor differences in the rate constants (39, 53-55). Absorption of blue-light (∼400-480 nm) by the embedded FMN triggers the formation of the singlet excited state almost instantaneously. The singlet decays to the triplet state in ∼2-4 ns via intersystem crossing. Finally, the triplet decays into the adduct state on time-scales of 0.2 to 10 μs. The slow accumulation of lit2 in LOV versions carrying the reactive cysteine is compatible with a short-lived triplet state and the low quantum yield of FMN reduction to hydroquinone (56). On the other hand, the rapid formation of lit2 state suggests that it originates from the long-lived triplet state (∼1 ms) (55), most likely preceded by a short-lived NSQ state. Interestingly, the isolated LOV domain of EL222 converts to the lit2 state much faster than the two-domain EL222 protein. This result suggests that the HTH domain stabilizes somehow the lit1 state relative to lit2. Accumulated evidence suggests that the reduction and protonation state of FMN, rather than the presence of a covalent bond between FMN and the LOV core, is the main determinant for all subsequent conformational changes (38, 42). Flavoproteins in general are sensitive to the redox state of their bound flavin (57, 58), and LOV proteins in particular may play a role in redox sensing in the cell (59, 60). The main structural changes in LOV proteins occur concomitantly with adduct formation (39, 54, 61). Such changes depend on the modules accompanying the LOV domain. For instance, large-scale structural changes do not occur during activation of the blue-light receptor YtvA,(62), while unfolding of the helices flanking the core LOV domain, A’α at the N-terminus and Jα at the C-terminus, has been reported for aureochromes (63, 64) and truncated versions of Avena sativa phototropin 1 (65-67). Photoinduced unfolding of α-helices has been observed for EL222 but so far has not been unambiguously assigned to a particular region (15). Contrary to FMN, which is essential for signaling, conserved residues in the LOV domain are not critical unless they affect FMN binding. Cysteine-less LOV variants can still transduce signals, albeit with lower efficiency and significant production of potentially damaging ROS (42, 68). A conserved glutamine residue appears to sense the protonation state at N5 via rotation of its side-chain, a movement that provokes long-range changes in the LOV domain (36, 54, 66, 69). However, glutamine-less variants can still be physiologically active when illuminated, suggesting that the coupling between FMN photochemistry and LOV holoprotein activation is more complex than previously predicted and likely proceeds via multiple pathways (70). Our results lend support to the view that the dynamic behavior of EL222 changes significantly after illumination and this shifts the conformational (domain separation, secondary structure) and oligomerization equilibria. Transcription factors can bind DNA cooperatively thereby strengthening their affinity (71, 72). Here we show that the DNA binding activity of EL222 is not an infinitely cooperative process, with no DNA interaction in the dark and complete association under light conditions. Rather, the different redox/adduct states of FMN in the dark, lit1 and lit2 states, differ in their dimerization (or more generally, oligomerization) properties, likely because of an altered propensity to populate an “open” conformation. Specifically, the lit1 state has the highest structural plasticity, i.e. tendency to change the secondary structure and oligomerize, and this may explain the increased DNA binding ability. In the presence of DNA, an EL222-DNA heterocomplex, predominantly of 1:1 stoichiometry, can be assembled, even in the dark. The complex assembly and disassembly kinetics, however, critically depend on the redox state of the embedded FMN chromophore, and thus on light. A stable 2:1 (protein:DNA) complex formation is most efficient in the lit1 state that is characterized by N_5_-protonated FMN and cysteinyl adduct formation. At this point, we can only speculate about the transcriptional activity of the different states. We expect that the encounter complex seen in the dark will have little ability to activate gene expression but may be important to determine the basal levels of transcription. Regarding the lit2 state, we hypothesize that it could probably sustain gene transcription, although less efficiently than the lit1 state. Since increasing the intensity of light is one way to achieve higher transcriptional output from optogenetic systems (8-10, 48), our findings may be taken into account to design more efficient EL222-inducible systems with improved dynamic ranges.

Our results highlight the complementarity between different structural and biophysical methods. MX delivers near-atomic-resolution structural information in the crystalline state but it is unclear to what extent restrictions imposed by crystal packing are affecting EL222 conformational changes (73). NMR is ideally suited to identify conformational subpopulations and their exchange kinetics in solution (74). It also provides atomic resolution structural and dynamical information, as long as the particle sizes are small enough. IR spectroscopy is sensitive to protein secondary structure changes and oligomerization, also in solution, but without any size limitation. However, IR data are difficult to interpret in the absence of additional information (31). UV/Visible spectroscopy is an excellent choice to monitor structural and electronic changes in FMN, including the formation of an FMN-cysteine adduct, but provides virtually no information beyond the chromophore. Therefore, only the combination of these different techniques allowed us to derive a comprehensive picture of the EL222 photocycle in the absence and presence of DNA.

## Materials and Methods

### Protein and DNA preparation

Five versions of EL222 were prepared (see **Table S5**). Most studies employed the EL222(17-225) variant (following UniprotKB Q2NB98) numbering, which differs from PDB numbering by an interval of -3). This is a quasi-full-length variant, referred to as EL222-WT, lacking part of the disordered N-terminus known to promote protein instability.(6) We also prepared a mutant lacking the conserved cysteine (EL222-C78A),(55) a slow recovery quadruple mutant (EL222-AQTRIP),(18) and a variant with a non-canonical 4-cyanophenylalanine (CNF) residue at position 31 (EL222-W31CNF).(15) We also prepared a shorter version, EL222-LOV containing only the LOV domain, i.e., lacking the linker and HTH domains. The purification tags were removed as previously described for EL222(17-225).(15) The final proteins contain only native residues except for the C-terminal alanine “scar” and the GEF sequence at the N-terminus of EL222-LOV. All protein handling steps were done under dim light. Proteins were flashed-frozen in liquid nitrogen and stored at -80°C until used.

### Mass spectrometry

All proteins were analyzed by mass spectrometry to determine the molecular weight (**Fig. S17**). Proteins were diluted with 100 μL of 5% acetic acid in water and loaded onto Opti-trap C4 cartridge (Optimize Technologies), washed 4× with 250 μL of 5% acetic acid in water and eluted with 100 μL of 80% acetonitrile, 5% acetic acid. Proteins were analyzed by direct infusion using syringe pump at a flow rate 2 μL/min connected with an electrospray ion source of 15 T solariX XR FT-ICR mass spectrometer (Bruker Daltonics). The mass spectrometer was externally calibrated using 1% (w/w) sodium trifluoracetate. Proteins were measured in positive mode with 2 M data acquisition. The data were processed using SNAP algorithm, a part of DataAnalysis 4.4 software (Bruker Daltonics).

To determine post-translational modifications in the EL222-WT batch used crystallization, the protein was digested using trypsin and further analyzed by LC-MS/MS using a timeToF Pro mass spectrometer (Bruker, Germany).

Forward and reverse DNA oligonucleotides (**Table S5**) were purchased separately (Sigma), dissolved in MES 50 mM pH =6.8 NaCl 100 mM, mixed in equimolar ratios and annealed by first heating up to 95 °C and then cooling down to room temperature (1 °C/min). The resulting double-stranded DNA was kept at 4 °C and used within one week.

### NMR spectroscopy

All NMR experiments were performed at 25°C on Bruker Avance III HD spectrometers (700, 850 or 950 MHz ^1^H frequency) equipped with helium-cooled triple-resonance probes (HCN TCI 5 mm) and pulsed z-field gradient. EL222 samples were prepared as 300 μL protein solutions at 200 μM concentration (unless otherwise stated) dissolved in 25 mM MES buffer, pH 6,8, 50 mM NaCl (plus the addition of 5% deuterium oxide). Samples were placed in NMR Shigemi tubes of 2 cm length and 5 mm diameter. ^1^H-^15^N correlation spectra and 3D NMR experiments for backbone assignments (HNCO, HNCACO, HNCA and HNCOCA) were obtained using Band-selective Excitation Short-Transient Transverse Relaxation-Optimized Spectroscopy (BEST-TROSY) (75) correlation experiments, as implemented in the NMRlib sequence library package (76) (which can be freely downloaded from the IBS website (http://www.ibs.fr/research/scientific-output/software/pulse-sequence-tools). Customized H(N)C, H(NCO)N and H(N)CO experiments were used for NMR resonance assignments of the FMN moiety in the different photo-stationary states. Translational diffusion of the protein and protein:DNA complexes in solution were measured by 1D ^1^H Diffusion Ordered Spectroscopy (DOSY) experiments (77) .focusing on the amide or methyl ^1^H spectral regions. A series of 1D spectra were recorded with varying gradient strength and a total acquisition time of approximately 20 min. ^15^N T_2_ transverse relaxation experiments were performed on uniformly ^15^N-^13^C-labelled protein samples using standard NMR pulse sequences (78). The NMR signal decay was sampled for 10 relaxation time points varying between 0 and 128 ms. The rotational diffusion constants of proteins and protein/DNA complexes, related to their particle size, has been determined by TRACT experiments (79), which measures the relaxation rate difference of the two doublet components of a ^1^H-coupled ^15^N (amide group) in a series of 1D ^1^H experiments.

The recorded NMR data were processed and analyzed using Bruker Topspin 3.5 and CCPNMR V2 software (Collaborative Computing Project for NMR; https://ccpn.ac.uk/software/version-2/version-2-downloads/) In-situ NMR sample illumination has been achieved as described previously (80). A laser source of a typical power of 70 mW, emitting maximally at 488 nm, was used to photoconvert the samples. This corresponds approximately to a power density of 260 mW/cm^2^ at the top of the NMR sample tube.

### IR spectroscopy

Steady-state infrared spectra of EL222 variants were measured with a Bruker Vertex 70v FTIR (Fourier-transform infrared) spectrometer equipped with a globar source, KBr beamsplitter and a liquid nitrogen cooled mercury cadmium telluride (MCT) detector. The protein samples in MES 50 mM NaCL 100 mM pH=6.8 were centrifuged at 18000 g for 15 minutes to remove any precipitants before measurements. For measurements in the Amide I band (EL222-WT and EL222-C78A samples), D2O was employed, while for measurements in the “transparent window” (sulfhydryl band of EL222-WT and EL222-C78A, and nitrile band of EL222-W31CNF), H_2_O buffer was utilized instead. D_2_O buffer exchange was done by a centrifugal concentration method using 10 kDa cutoff filters. The aperture value was set in order to get maximum signal without saturating the detector. Circa twenty-five μl of sample was loaded into a temperature controllable demountable liquid cell with CaF_2_ windows and a 50 μm Teflon spacer. Eighteen spectra were recorded at 20 °C and 15-minutes intervals. The initial 14 spectra were recorded while the protein was illuminated to induce the lit state, achieved using two LEDs emitting a power of 100 mW/cm^2^ onto the sample throughout the measurement. The final four measurements were taken with the lights turned off. All difference spectra were recorded against the initial spectra in the absence of illumination (dark state). The double difference spectra of **Fig. 6c** were calculated as (lit_x_-dark)^*DNA*^ minus (litx-dark)^*free*^, where the subindex x can be 1 or 2 and *DNA/free* denotes the presence or absence of DNA, respectively. Protein samples had concentrations between 0.5 and 1 mM. In the case of protein-DNA mixtures equimolar ratios were employed. A water vapor spectrum was measured with the same parameters and utilized to remove any water vapor contamination present in the obtained protein spectra. The spectra underwent baseline correction using OriginPro (OriginLab).

### UV/Visible spectroscopy

The UV-Visible absorbance spectra of the EL222 protein within the 320-550nm range were obtained using a Specord 50 Plus spectrometer (Analytik Jena). Measurements were conducted at 20°C in a quartz cuvette (Hellma Analytics) with a 10mm path length for quantification purposes and a 1mm path length for lit state induced measurements, with sample volumes of 70 μl and 600μl respectively. To ensure an airtight environment for lit state measurements, the cuvette opening was sealed with a PTFE lid and parafilm. Prior to dark state sample measurements, MES buffer served as the background measurement. The protein’s lit state was induced by three hours of continuous illumination using three 455-nm LED with a power of 1 mW/mm^2^, covering the entire cuvette. Subsequently, the LEDs were deactivated, and spectra were recorded.

## Data availability

X-ray structures have been deposited in the Protein Data Bank under accession codes 8A5R (EL222 dark) and 8A5S (mix of EL222 dark and EL222 lit1). NMR chemical shifts have been deposited in the Biological Magnetic Resonance Data Bank under accession codes BMRB 52621 (EL222 dark), BMRB 52620 (EL222 lit1), and BMRB 52622 (EL222 lit2). All input files required for classical MD and metaD simulations for the *oxi, cov*, and *red* models in GROMACS, including configuration files, COLVAR and HILLS files, and post-processing scripts, are provided in Zenodo repository, accessible at https://doi.org/10.5281/zenodo.13889603. All infrared and UV/Vis spectra can be found in the same Zenodo link.

## Acknowledgments

The work was supported by the project Structural dynamics of biomolecular systems (ELIBIO) (CZ.02.1.01/0.0/0.0/15_003/0000447) from the European Regional Development Fund (ERDF) and the Ministry of Education, Youth and Sports (MEYS) of the Czech Republic, and by the project “Time-resolved vibrational spectroscopy of proteins assisted by genetically encoded non-canonical amino acids” from the Czech Science Foundation (24-11819S). The Institute of Biotechnology of the Czech Academy of Sciences acknowledges the institutional grant RVO86652036. We acknowledge CF Biophysics, CF SMS, CF Crystallization, CF Diffraction of CIISB, Instruct-CZ Centre BIOCEV, supported by MEYS CR (LM2023042) and ERDF-Project “UP CIISB” (No. CZ.02.1.01/0.0/0.0/18_046/0015974). Financial support from iNEXT-Discovery (871037) funded by the Horizon 2020 program of the European Commission, and the IR INFRANALYTICS (FR2054) is acknowledged. This work used the platforms of the Grenoble Instruct-ERIC centre (ISBG; UAR 3518 CNRS-CEA-UGA-EMBL) within the Grenoble Partnership for Structural Biology (PSB), supported by FRISBI (ANR-10-INBS-0005-02) and GRAL, financed within the University Grenoble Alpes graduate school (Ecoles Universitaires de Recherche) CBH-EUR-GS (ANR-17-EURE-0003). IBS acknowledges integration into the Interdisciplinary Research Institute of Grenoble (IRIG, CEA). Computational resources were supplied by the project “e-Infrastruktura CZ” (e-INFRA ID: 90254) provided within the program Projects of Large Research, Development and Innovations Infrastructures and with the support of ELIXIR CZ Research Infrastructure (ID LM2023055, MEYS CR). We thank Dr. Kevin Gardner (CUNY Advanced Science Research Center, New York, USA) for fruitful discussions and critical reading of the manuscript.

