## Supplementary information for "Light-dependent flavin redox and adduct states control the conformation and DNA binding activity of the transcription factor EL222"

**This PDF file includes:**

Supporting text (Notes S1 to S3)
Figures S1 to S17
Tables S1 to S5
Legend for Movie S1
SI References

**Other supporting materials for this manuscript include the following:**

Movie S1

**References** ........ **29-31**

### Supporting Information Text

#### Note S1: X-ray crystallography experiments.

Protein crystallization.

EL222-WT at 10 mg/ml concentration in a buffer containing 25 mM PIPES, pH 7.2, with 65 mM NaCl was used for crystallization. Several commercial crystallization screens were used for initial screening using the vapour diffusion method and the sitting drop setup. The final optimized reservoir solution (0.11 M MES, pH 6.5, 0.22 M $MgCl_2$, 21% (w/v) PEG 8000) was based on condition D9 from the XP screen (Jena Bioscience, Germany). Crystals grew in the sitting drop setup using CrystalQuick™ 96-Well Protein Crystallization Plates (Greiner Bio-One, Austria), 50 μl of reservoir solution and drops composed of 0.8 μl of protein solution and 0.4 μl of reservoir solution. Crystals grew in two months at 18 °C.

X-ray diffraction data collection and processing.

Crystals of both EL222dark and EL222light were retrieved from the same drop and soaked in a solution containing 80% (v/v) of the reservoir solution and 20% (v/v) glycerol for 60 s. Then crystals were mounted using MicroMeshes™ 400/25 (MiTeGen, USA) on the diffractometer with a humidity control system HC-Lab (Arinax, France) under the stream of air with 96% relative humidity at room temperature. Crystals were conditioned for 30 min, letting the crystal equilibrate to air with pre-set humidity, and then vitrified in a cold nitrogen gas stream (100 K). All manipulation of the crystal for the structure EL222dark was done under low ambient light. For the structure EL222light, the crystal was illuminated using a fiber-coupled LED light source (M470F3 Thorlabs, USA) and a 1 mm fiber patch cable (M59L01, Thorlabs, USA) producing 30 mW of power (at 1 cm distance between the end of the cable and the crystal) of 470 nm wavelength light during conditioning and data collection. X-ray diffraction data were collected from both crystals at 100 K using a D8 Venture diffractometer with a Photon II detector (Bruker, Germany) and a MetalJet D2 X-ray source (Excillum, Sweden). Data for EL222dark were processed using the Proteum3 software package (Bruker, Germany), and for EL222light using XDS (1). Aimless (2) was used for analysis of both datasets and for EL222 light also for data merging. Identical sets of reflections were selected for calculation of $R_{free}$ for these two structures. Data collection and processing statistics are reported in **Table S1**.

Structure determination and refinement.

For both structures phases were obtained by molecular replacement using Molrep (3) and a structure of identical protein (in the dark state) as a template (PDB ID: 3P7N (4)). Both structures were manually built/modified using Coot (5) and O (6)[ref], refined using REFMAC5 (7) with calculation of $R_{free}$ as a cross-validation method and with hydrogens in riding positions. Structures were validated using the PDB validation service (8). Composite omit electron density maps calculated using Phenix (9) were used in building and validation of both structures. Flavin mononucleotide (FMN) in the light state in EL222light was modelled covalently bonded to Cys78 with 30% occupancy. Its geometry was modelled in accordance with similar structures in the light state containing the same FMN – cysteine covalent link (PDB ID: 1JNU,(10) 2V1B, 2V0W,(11) 2PR6,(12) 6S46 (13)) and refined using specific restraints defined for this purpose based on these structures. The refinement statistics are reported in **Table S1**.

**Note S2: Lit1-to-lit2 photoconversion kinetics.**

NMR.

In the case of NMR experiments, the decay of lit1 population, which coincides with the built-up of lit2 species, was fitted with an exponential function.

IR.

In the case of infrared spectroscopy, the time-resolved difference spectra (lights on minus light off), which do not directly report on the population of different species, were subjected to lifetime distribution analysis using the maximum entropy method as previously described. The spectra were first converted from the time domain to the lifetime domain using the inverse Laplace transform (14-16). Then the average dynamical content (*D*) (17, 18) was calculated by averaging over the whole wavenumber range. These ranges were 1600-1750 $cm^{-1}$, 2500-2650 $cm^{-1}$, 2220-2250 $cm^{-1}$ for the Amide I' (C=O stretching from protein and FMN), sulfhydryl stretching, and nitrile stretching regions, respectively. The peak maximum of the *D* lifetime distribution was taken as the lifetime of lit1 species.

The fitted lifetimes can be found in **Table S3**.

**Note S3: Computer simulations.**

System preparation.

The initial configurations for EL222 in *oxi*, *cov*, and *red* states of FMN (**Fig. 2d**), intended to emulate the experimentally observed dark, lit1, and lit2 states, were derived from crystal structures of dark state (PDB code: 8A5R, 1.85 Å, for *oxi* and *red*) and light state (PDB code: 8A5S, 1.85 Å, under 470 nm illumination, for *cov*). For *oxi* and *red* models, the FMN structure was extracted from protein-FMN complex PDB file and saved in PDB format for further processing. For the FMN-cysteine adduct formation in *cov* model, the side chain of the cysteine residue (Cys78 residue in EL222 protein) was retained as part of the FMN structure for the ligand parametrization step. The protonation states of histidine residues in the protein structures were assigned based on the hydrogen bond network and solvent accessibility, simulating physiological conditions (pH = 7.4). All hydrogen atoms in the protein structure were added using tleap, integrated in AmberTools (19). The protonation states of the FMN ligand were modeled using Chimera software, guided by experimental observations (20) (**Table S4**). Therefore, total charge of -2 has been assigned to FMN ligand in *oxi* and *cov* states and total charge of -3 has been assigned on *red* state due to N1-atom charged equilibrium preferences at physiological conditions (**Fig. S6**). These structures were optimized at DFT-D3/TPSS/def2-SVP level with the water-COSMO model for solvation treatment using ORCA 5.0.1 (21).Topology and parameter files for the FMN ligand in *oxi*, *cov* and *red* states were generated using the General Atom Force Field (GAFF) in ANTECHAMBER module of AmberTools, creating .prep and .frcmod files (22, 23). Missing parameters not defined in GAFF were added using the parmchk2 tool.

In *oxi* and *red* states, where the FMN ligand is docked into EL222 binding pocket without covalent bond formation, ligand and protein parametrization follow standard protocols in Antechamber and tleap modules of AmberTools. For FMN-cysteine adduct model in *cov* state, after creating .prep and .frcmod files using FMN-cysteine adduct structure using ANTECHAMBER, cysteine side chain's atoms were removed from .prep file and their corresponding atomic charges were redistributed to ensure a total charge is an integer in modified .prep file. Additionally, residue name for Cys78 residue in protein PDB file was changed from CYS to CYB. After preparing protein and FMN structures and their parameter files for investigated models, tleap from AmberTools was used to merge these structures and form a covalent bond between the C4a atom of the FMN moiety and the sulfur atom of CYB residue in the protein (bond 78.SG 207.C4A command). This required loading force field parameter files for the protein (Amber force field, ff14SB (24), and the ligand (prepared .prep/.frcmod files in GAFF) and their structures. Due to application of two different force fields for the FMN ligand and protein residues in the *cov* complex, atom types for C4a-S covalent bond and angles around this covalent bond in CYB.prep and CYB.frcmod files were updated accordingly. The final molecular models of EL222 in *oxi, cov*, and *red* states of FMN, which were used as starting structures for all subsequent simulations, are depicted in Fig. 4A).

Given the known involvement of glutamine (Gln) residue rotation in the conformational dynamics of other photosensitive proteins, particularly within the LOV-FMN binding domain, it is essential to investigate whether a similar behavior occurs in the LOV domain of our EL222 protein model upon light exposure (for Gln141 residue, details are given in Figure 1 and Fig. S2). To explore this, we conducted two series of molecular dynamics (MD) simulations with initial structures taken from PDB ID: 8A5S, each with a cumulative runtime of 3 μs for *cov* state (*cov*$^{MD}$): Position (a) one where the Gln141/Nε2 formed a hydrogen bond with FMN-N5, as observed in lit structure of LOV-binding domain, and alter position (b) a second where the Gln141/Oε1 formed the hydrogen bond with FMN-N5. We assessed the conformational behavior of Gln141 residue throughout these MD simulations by measuring the distance between the donor and acceptor hydrogen-bond atoms in the FMN chromophore and Gln141 residue (i.e.; Gln141/Oε1 with FMN/N5, Gln141/Nε2 with FMN/N5 and Gln141/Nε2 with FMN/O4 distances). In model (a), spontaneous rotation of the Gln141 side chain was observed for 30% of the simulation time. In contrast, the Gln141/Oε1 configuration in model (b) remained stable throughout whole simulation time, with no reversion to the non-rotated or Gln141/Nε2 form, even in the absence of external force. While the rotation of the Gln141 side chain was favored in *cov* state, it was not stable in *red* state (*red*$^{MD}$). Therefore, the Gln141 side chain was left in its original orientation, as observed in *oxi* state (*oxi*$^{MD}$) or in X-ray dark structure for *red* state (*red*$^{MD}$).

Molecular dynamics (MD) simulations.

Since GROMACS package was used for simulation step in this study, ParmEd tool (25) was used to convert .prmtop and .inpcrd files prepared in AMBER for *oxi*, *cov* and *red* to .top and .gro GROMACS compatible files. Each protein-FMN complex was placed in a truncated dodecahedron box and solvated with TIP3P water molecules to a depth of at least 11 Å (gmx editconf and gmx solvate modules). The solute was neutralized with sodium chloride ions to reach a physiological salt concentration of 0.1 M (gmx genion). All-atom molecular dynamics (MD) simulations were performed using the GROMACS package version 2023.2 on GPU hardware (26, 27).

Energy minimization was conducted using the steepest descent algorithm (50000 steps) to optimize the initial structure, with a step size of 0.01 nm for each minimization step. The particle mesh Ewald method (PME) was employed for long-range electrostatic interactions with a cut-off 10 Å for non-bonded interactions (van der Waals and direct Coulomb). After energy minimization, the system was equilibrated under NVT condition using V-rescale modified Berendsen thermostat at 300 K (28) (50000 steps with a time step of 2 fs). Equilibration of pressure is conducted under NPT ensemble condition with Parrinello−Rahman barostat (29) (50000 steps with a time step of 2 fs). Position restraints were applied to protein to prevent significant deviations from the initial structure. The LINCS algorithm was used to restrain bond lengths. *Oxi*, *cov*, and *red* states were subjected to 3 independent MD simulation replicas, each lasting 1 microsecond (3 microseconds cumulative MD). Protein stability during simulation was evaluated by root-mean-square deviations (RMSD) considering backbone atoms in $oxi^{MD}$, $cov^{MD}$, and $red^{MD}$ simulation runs. The fluctuation of protein residues for $oxi^{MD}$, $cov^{MD}$, and $red^{MD}$ at Cα-atom level was evaluated using the root mean square fluctuation calculation (given in **Fig. S13**) using Bio3D R (30).

<u>Metadynamics (metaD) simulations using PLUMED.</u>

Metadynamics simulation (metaD) is a powerful force sampling method that introduces a bias potential along selected collective variables (CV) in the free energy surface (FES) of a protein (31). Collective variables are specific parameters that summarize the relevant degrees of freedom of a system, such as distances, angles, or dihedrals, capturing important structural changes during the simulation. The bias potential in metaD prevents the system from revisiting previously Therefore, metaD accelerates the exploration of the conformational landscape in proteins compared to classical MD simulations, which often require extensive simulation time frames spanning microseconds.

To investigate the dissociation of the LOV and HTH domains in the different FMN states of EL222 protein (*oxi*, *cov*, and *red*), we employed Metadynamics-PLUMED simulations (32-34) in Gromacs (Gromacs version 2023.2-plumed_2.10.0_dev), using a single collective variable (35), starting from equilibrated MD structures (NPT step). This CV was defined as distance between center-of-mass (COM) of all residues within HTH-4α region (sequence of Ser213-Ala223 residues) and the COM of specific residues within the beta strands of the LOV domain (Pro103, Leu105, Glu107, Ala121, Leu123, Ala125, Leu138, Ser140, Val142 residues). An upper wall potential was applied to this distance (2.0 nm), with a specified threshold and force constant, to limit the maximum allowed separation. Metadynamics bias was applied every 1000 steps, with an upper wall constraint set at a distance of 20 Å and a spring constant (KAPPA) of 150.0 kJ/(mol*nm$^2$). The strength of this potential increases quadratically with a force constant of 150.0, ensuring that the system remains confined within a specific region of the collective variable space during the simulation. Gaussian hills with a height of 0.05 kJ/mol and a width of 0.25 nm were added to the bias potential. We performed 160 simulation replicas, each lasting 50 ns with a 2 fs time step. Distance and bias data were recorded every 5000 steps in the COLVAR and HILLS files.

The resulting COLVAR files, containing time-series data of the biased collective variable (including time, distance, and bias potential), for 160 simulation replicas related to $oxi^{metaD}$, $cov^{metaD}$ and $red^{metaD}$ models were collected and analyzed. To determine LOV-HTH domain separation time, we used a custom R script for each COLVAR file (**bias-distance-time-plots.R**, provided in supplementary materials deposited in the Zenodo repository). The primary objective of the script was to generate comparative plots for these variables across multiple simulation replicas. Plots of bias vs time, distance vs time, and diff-bias vs time for each COLVAR file were compiled into PDF files for further analysis. Additionally, plots of LOV-HTH separation distance over metaD simulations were generated using **LOV-HTH_separation_distance_PLUMED.tcl** script, are provided in Zenodo repository. The points at which the LOV-HTH domain separation occurs are indicated as vertical dashed lines in these plots (an example is given in **Fig. S14a**).

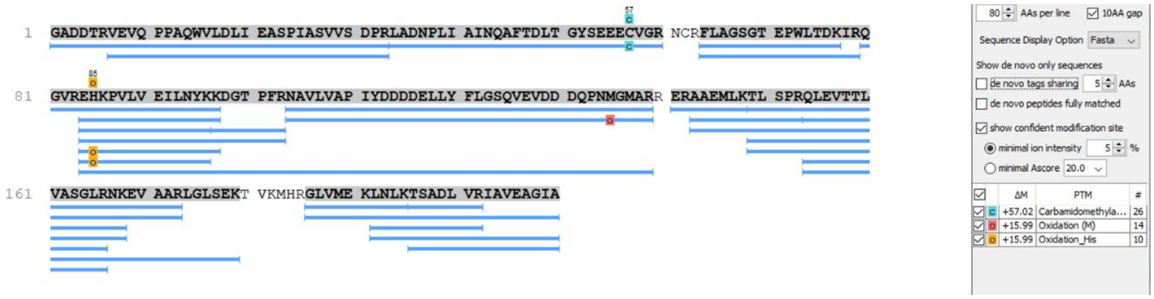

**Fig. S1. Mass spectrometry confirms the presence of chemical modifications in the EL222 sample used for crystallization.**

Tryptic digest of EL222 showing the sequence coverage and the presence of oxidized histidine at residue position 101.

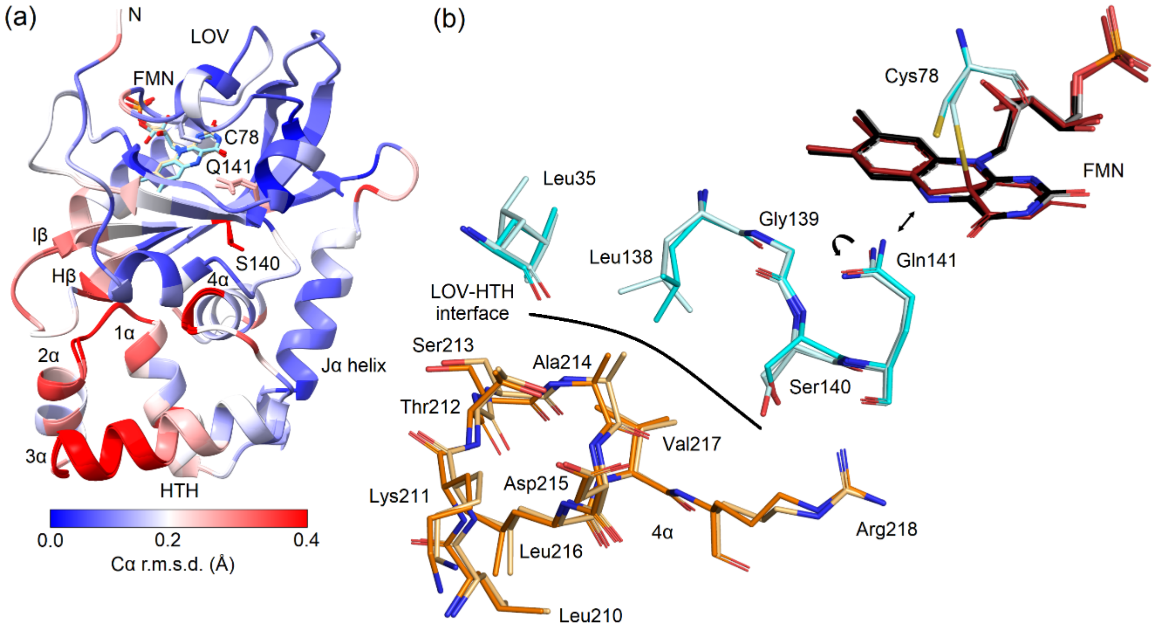

**Fig. S2. X-ray crystallography suggest the propagation of conformational changes in EL222.** (A) EL222 secondary structure representation colored by Cα r.m.s.d. between the dark and light structures, calculated using UCSF ChimeraX and the "matchmaker" command. The overall standard uncertainty of the positional parameters based on a maximum likelihood residual for EL222 structures obtained in the dark (PDB ID: 8A5R) and lit by blue light (PDB ID: 8A5S) are 0.072 Å and 0.079 Å, respectively. FMN, Cys78, Ser140 and Gln141 are shown as sticks. Selected elements of secondary structure are labelled. (B) Details of changes in molecular contacts on the LOV-HTH interface caused by the change in hydrogen bonding between the Gln141 side chain and the FMN-N5 atom (black double arrow), and consequential flip of the Gln141 side chain (black curved arrow). Selected residues of the LOV domain are shown as sticks, labelled and colored cyan/pale cyan for dark and lit structures, respectively. The residues at the beginning and end of a part of the HTH domain 4α helix (orange/pale orange for dark and lit structures, respectively) with the largest coordinate changes are labelled. FMN is shown as sticks with carbon atoms colored black (dark state) or red (lit state). Molecular graphics for panel A was created using UCSF ChimeraX (36), for panel B using PyMOL (37).

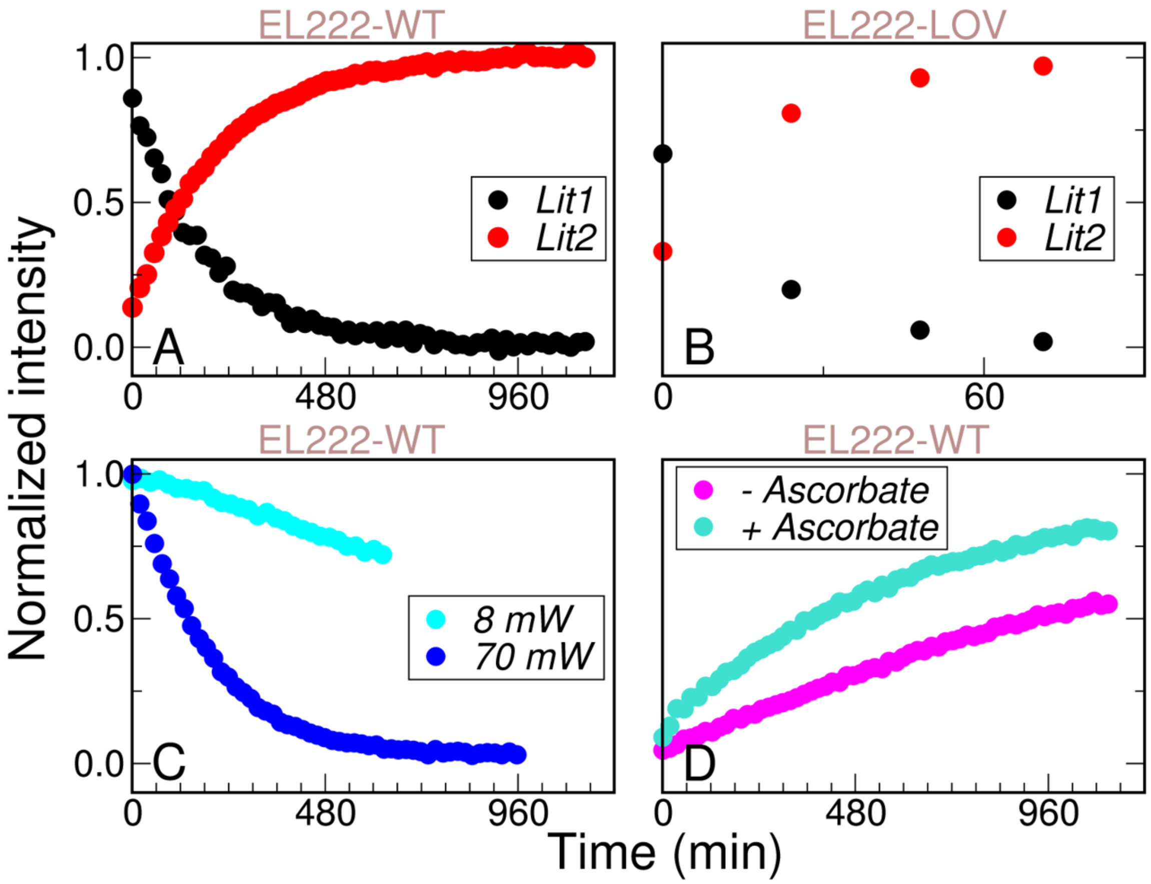

**Fig. S3. NMR-derived kinetics of lit1-to-lit2 conversion.** (a) EL222-WT kinetics (b) EL222-LOV kinetics. The normalized intensities were obtained as the sum of well resolved $^{1}H$-$^{15}N$ correlation peaks with different chemical shifts for the lit1 and lit2 states. (c) Similar data as shown in (a) for EL222-WT lit1 decay using different 405 nm illumination powers. A linear power dependence of the conversion kinetics on the light power is observed. (d) The conversion kinetics from lit1 to lit2 is redox-dependent: addition of a sacrificial electron donor (ascorbic acid) speeds up lit1-to-lit2 conversion kinetics.

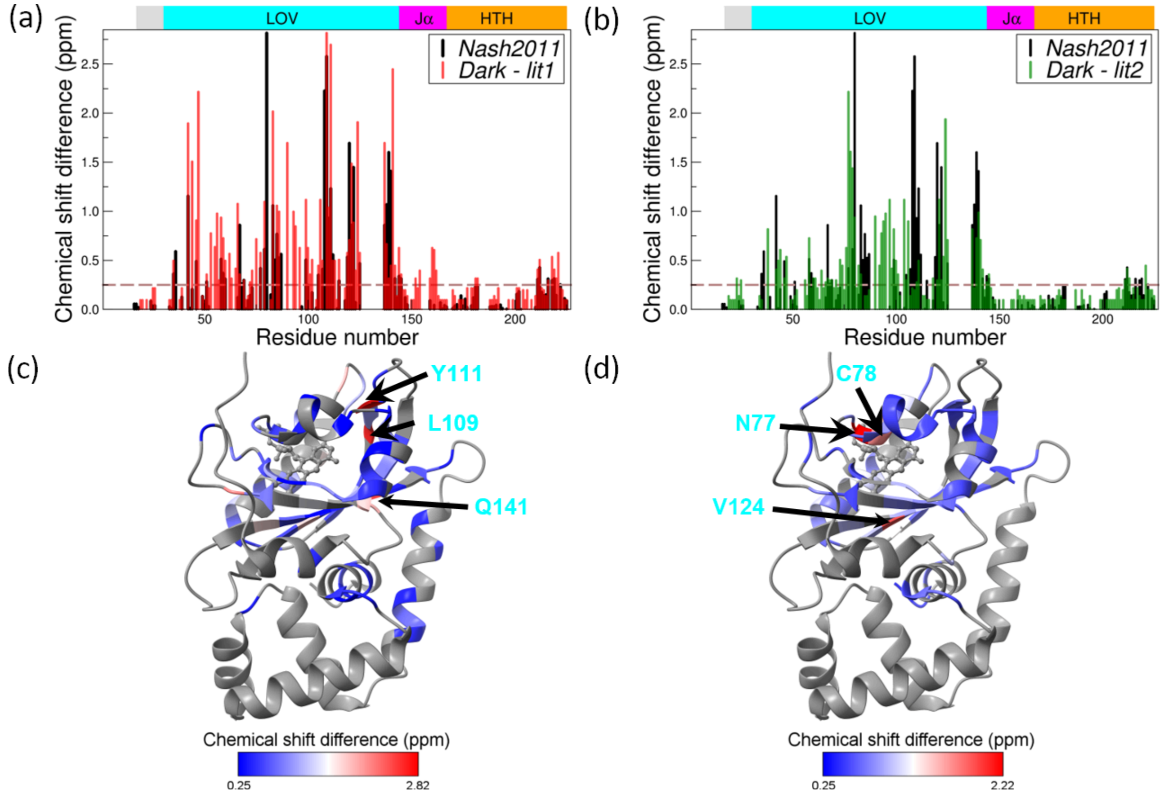

**Fig. S4. NMR chemical shift differences between the dark, lit1, and lit2 states of EL222.** $^{1}H$-$^{15}N$ chemical shift differences between (a) dark and lit1 states, and (b) dark and lit2 states. Scaled dark minus lit differences reported in Reference (6) of the main text are shown in black. Mapping these chemical shift differences onto the dark-state structure of EL222 (PDB ID: 8A5R) indicates that in both cases (c) lit1 and (d) lit2, conformational changes are localized mainly in the FMN-containing LOV domain, as well as the C-terminal end of the HTH motif. For the ease of visualization, a threshold of 0.25 ppm was applied. The three residues with the largest chemical shift differences are indicated.

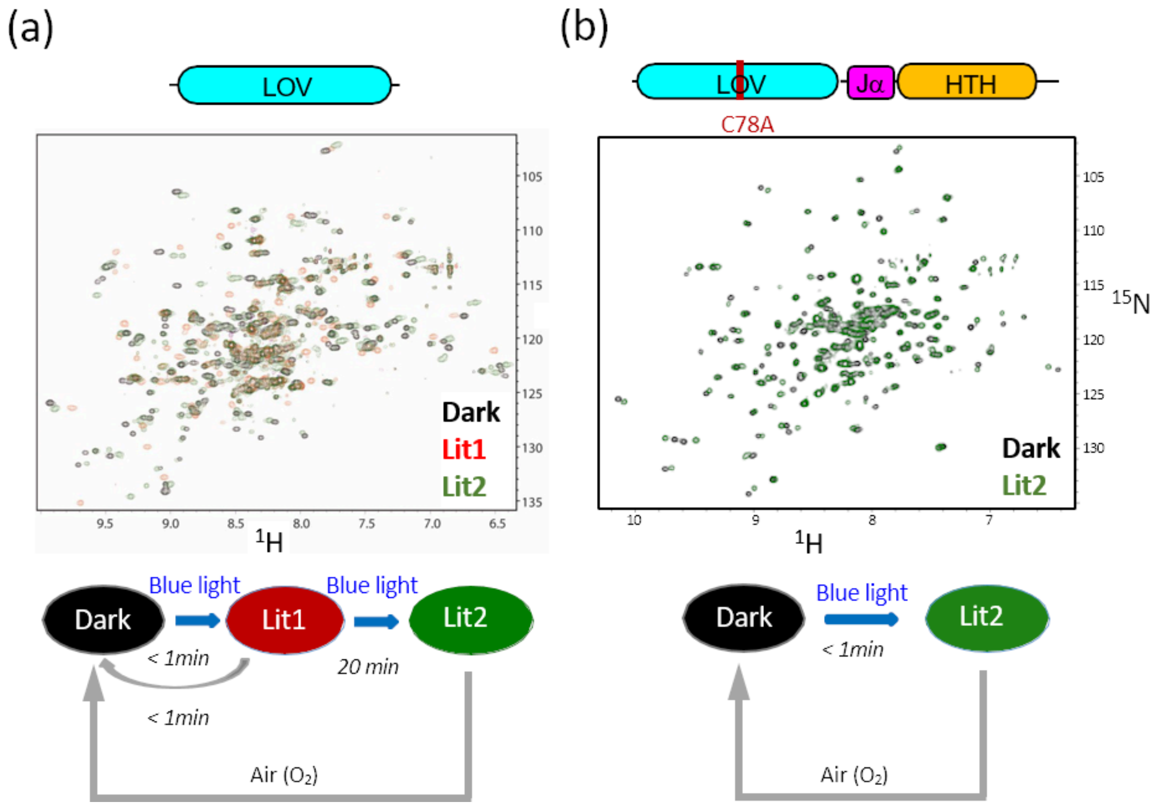

**Fig. S5. 2D-NMR spectra and proposed photocycle of two EL222 variants.** $^1$H-$^{15}$N correlation spectra (top) and scheme of the photocycle (bottom) of (a) EL222-LOV, and (b) EL222-C78A. Similar to EL222-WT, EL222-LOV shows three conformational states (dark, lit1, and lit2) with different conversion kinetics (lit1-to-lit2 transition is an order of magnitude faster in EL222-LOV relative to EL222-WT). EL222-C78A, which is devoid of the adduct forming cysteine residue, lacks the lit1 state and thus photoconverts directly to the lit2 state. All EL222 variants show the same oxygen dependence of the lit2-to-dark thermal recovery.

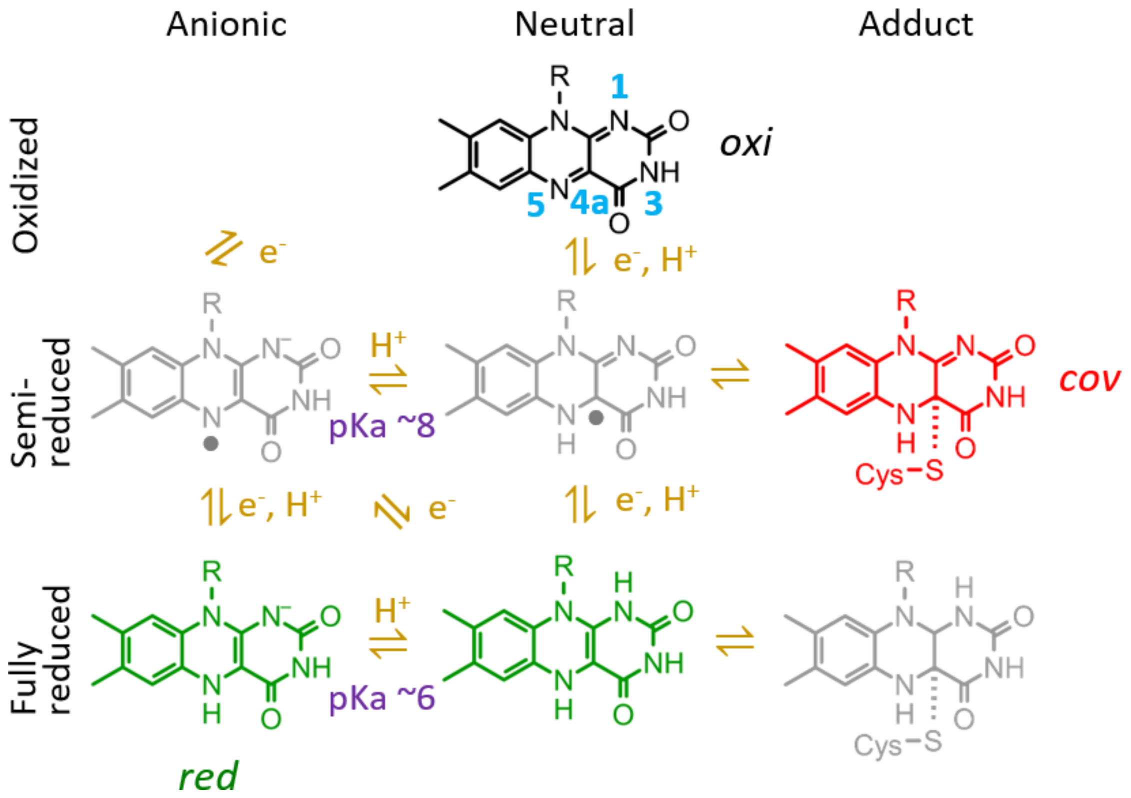

**Fig. S6. Major FMN species around neutral pH.** Isoalloxazine ring atom numbering is shown for oxidized flavin. The adjacent nearby cysteine (Cys78 in EL222) is indicated. Species colored in gray are possible intermediates in the LOV photocycle which, however, have not been detected under our experimental conditions. Species colored in black, red, and green, correspond to *oxi*, *cov,* and *red* FMN forms of **Fig. 2d**. Note that the neutral hydroquinone state of FMN-bound EL222 has only been detected at acidic pH (**Fig. S7a**, right panel).

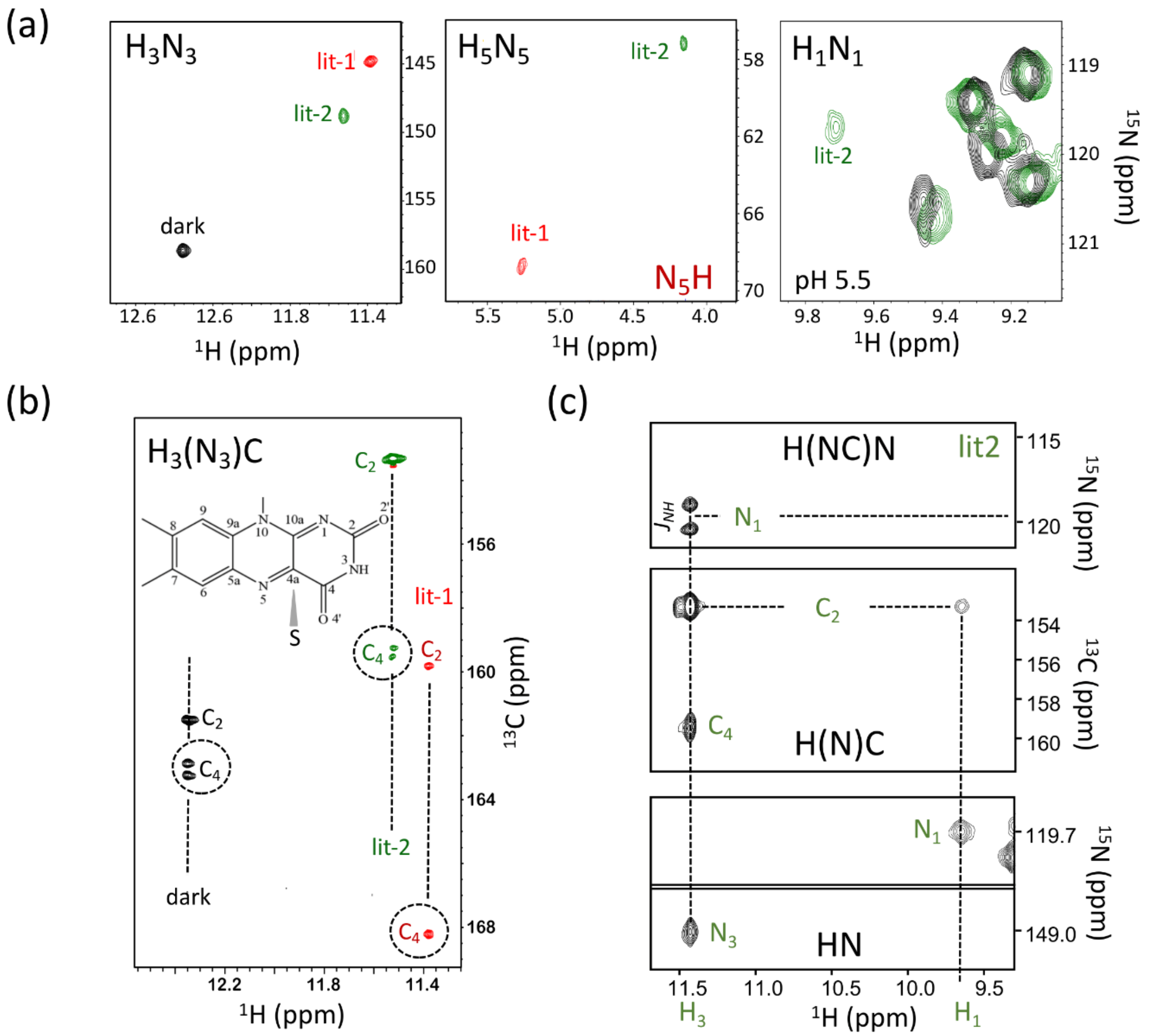

**Fig. S7. NMR determination of the redox and protonation states of FMN in the different EL222 states.** (a) Superposition of $^1H$-$^{15}N$ spectral regions of EL222 in dark (black contours), lit1 (red contours) and lit2 (green contours) states with HN groups from the FMN moiety assigned (labelled). Note that the $N_1H_1$ cross peak could only be detected under acidic conditions (pH 5.5) due to the low local pKa of this site. (b) NMR signature of adduct formation in lit1. An H-N-C experiment correlates the $H_3$-$N_3$ of FMN with the neighboring $C_2$ and $C_4$ carbons. Homonuclear $^{13}C$-$^{13}C$ spin decoupling over a bandwidth of 65 ± 15 ppm was applied during $^{13}C$ chemical shift labeling resulting in a single peak for the $H_3N_3C_4$ correlation peak (only) if the $^{13}C_{4a}$ chemical shift is within this bandwidth, which is expected in the case of adduct formation. (c) Additional HNC correlation experiments performed to confirm the assignment of the additional peak detected in the amide $^1H$-$^{15}N$ spectral region of the lit2 state (at acidic pH) to $N_1H_1$ (FMN).

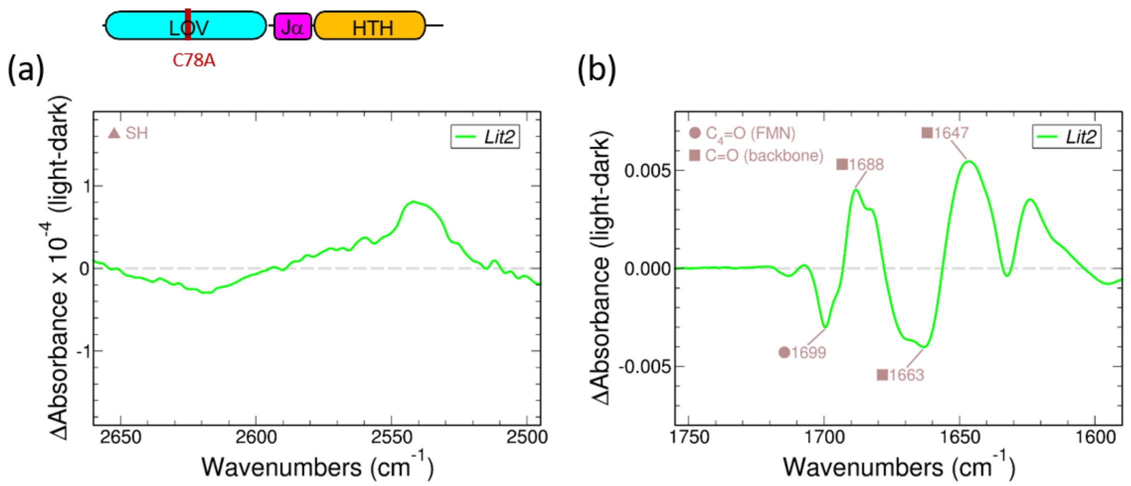

**Fig. S8. Infrared spectra of EL222-C78A.** (a) Lit-minus-dark difference IR spectra in the absorption region of SH bonds. (b) Lit-minus-dark difference IR spectra in the Amide I’ band.

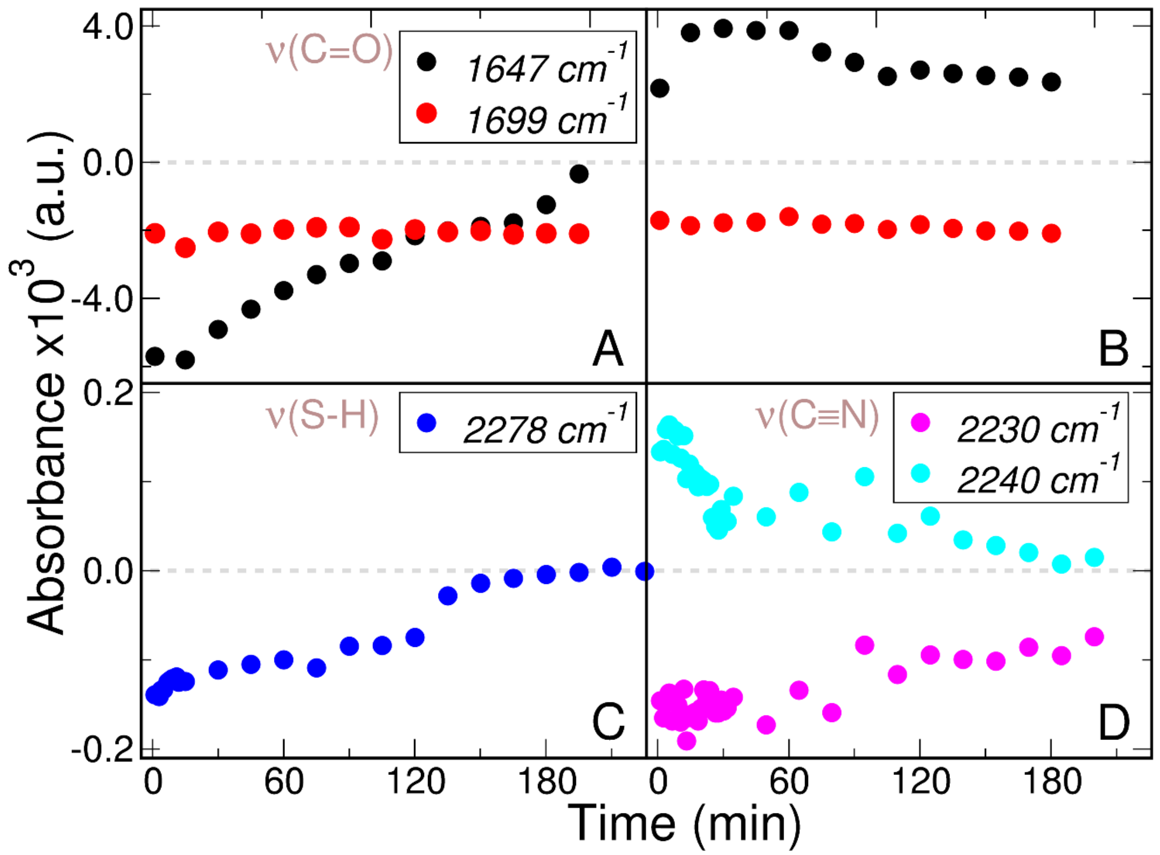

**Fig. S9. Infrared-derived kinetics under continuous illumination.** (a) EL222-WT in the Amide I' band. (b) EL222-C78A in the Amide I' band. (c) EL222-WT in the S-H absorption band. (d) EL222-W31CNF in C≡N absorption band. The tracked frequencies are indicated.

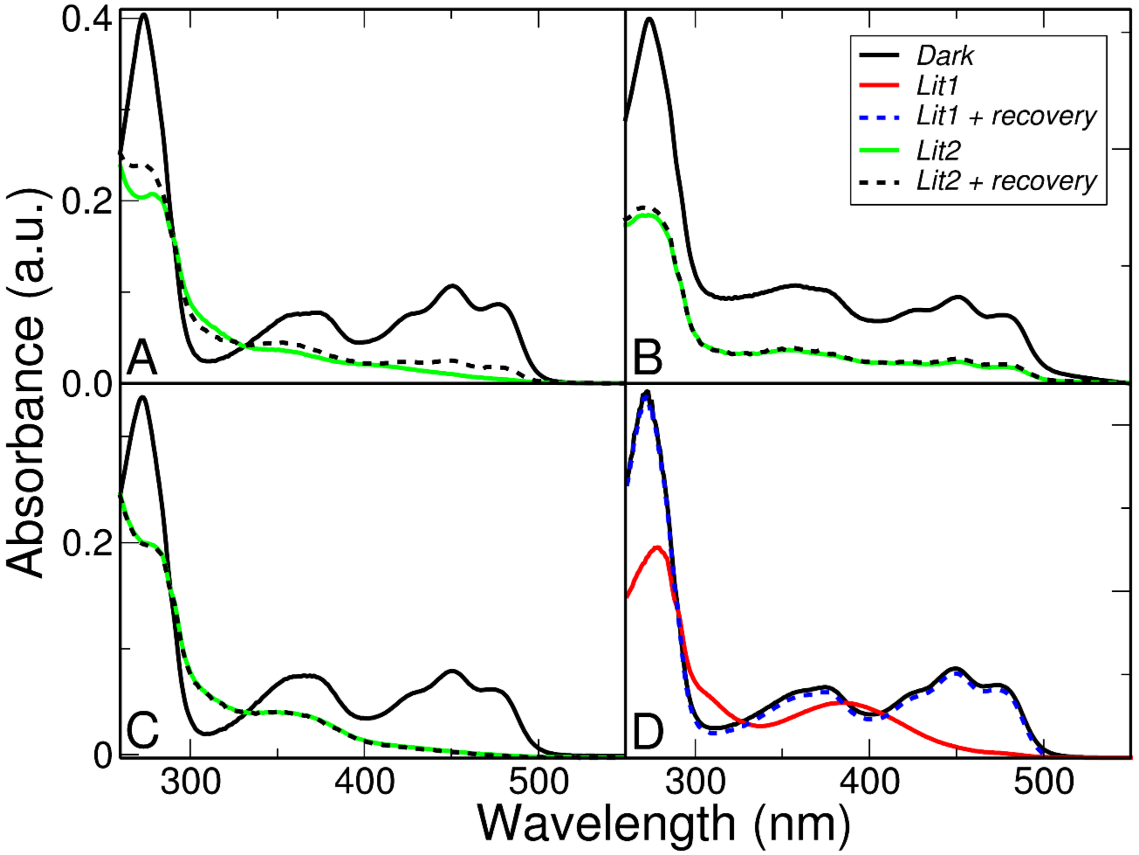

**Fig. S10. UV/Visible spectra of the studied EL222 variants along the photocycle.** (a) EL222-WT (b) EL222-WT at pH=5.5. (c) EL222-C78A. (d) EL222-AQTRIP. Spectra have been measured in MES 50 mM, pH=6.8 (except panel b), NaCl 100 mM. “Dark” spectra were recorded taken in the absence of illumination. “Lit1” spectra were taken after 1 min illumination. “Lit1 + recovery” spectra were taken after 1 min illumination and an additional hour in the dark. “Lit2” spectra were taken after 3 hours of continuous illumination. “Lit2 + recovery” were taken after 3 hours of continuous illumination and an additional hour in the dark.

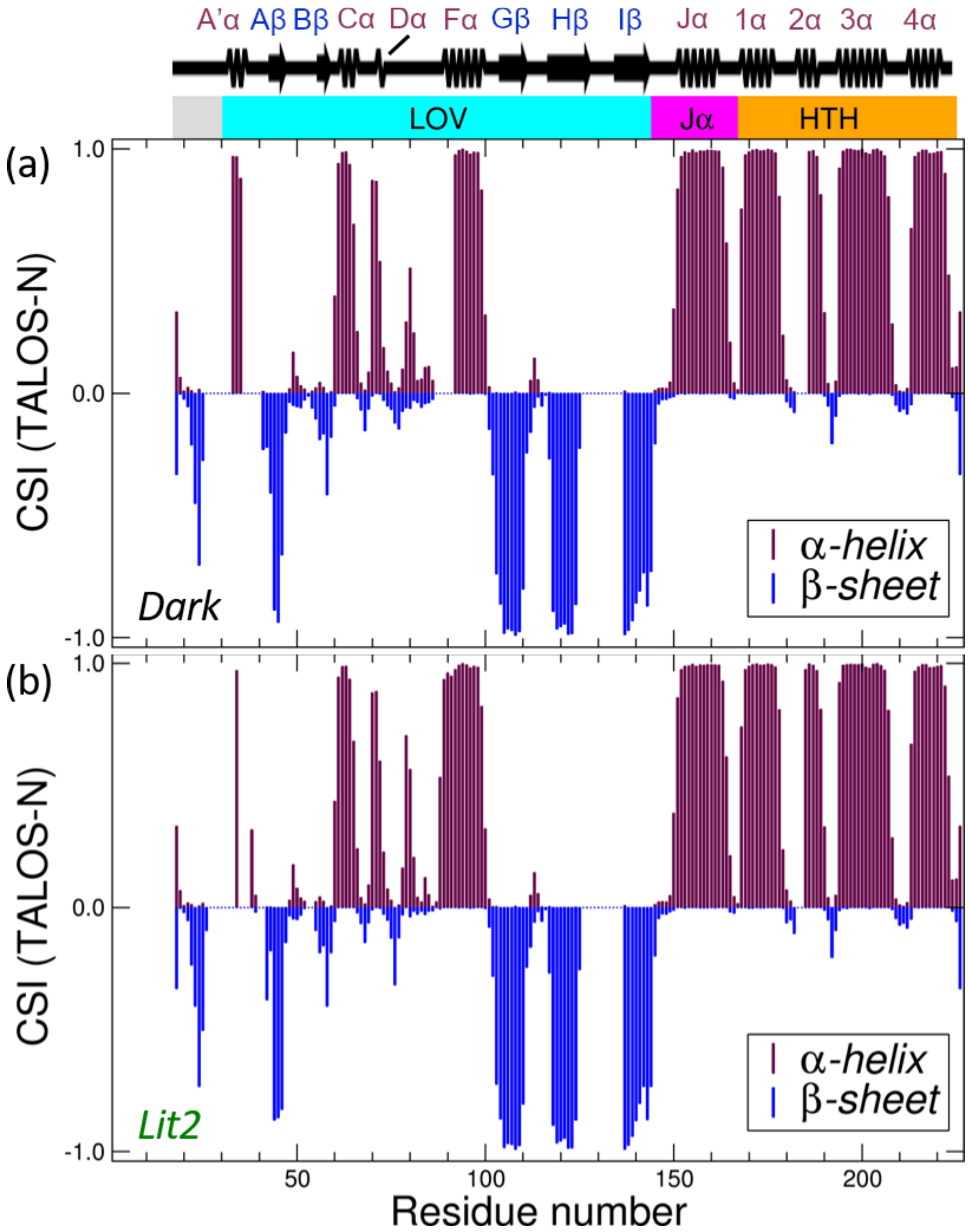

**Fig. S11. NMR-derived secondary structure content in EL222.** TALOS prediction (refs. 27 and 28 of main text) of EL222 secondary structure based on NMR chemical shifts (H, N, CO, CA) of dark (a) and lit2 (b) species. The secondary structure is essentially the same for the 2 states (similar $^{13}C$ chemical shifts) and in agreement with the secondary structure derived from our crystallographic model (8A5R, top panel). Secondary structure diagram has been generated with SSDraw (38).

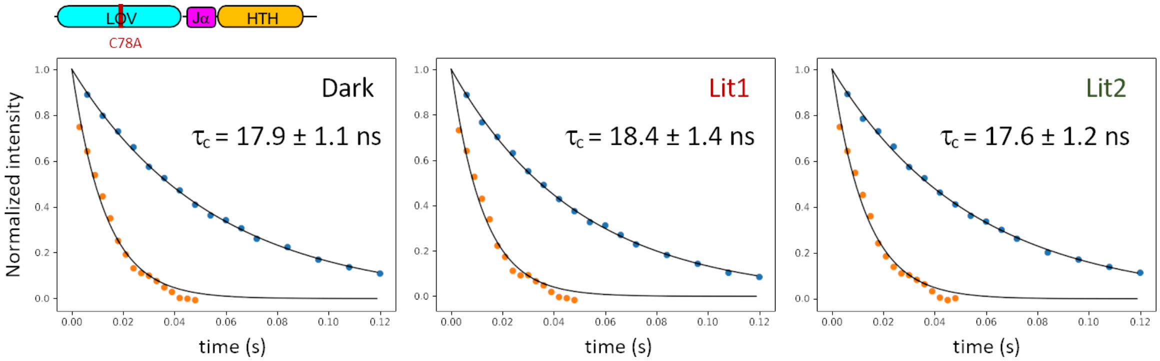

**Fig. S12. $^{15}$N TRACT measurements to determine the rotational tumbling correlation time of EL222-WT.** Dark (*left*), lit1 (*middle*), and lit2 (*right*) states. The normalized intensity of the two doublet components of a $^{1}$H-coupled $^{15}$N are plotted as function of a relaxation delay, and fit to an exponential function to derive the rotation tumbling correlation time $\tau_c$. No significant difference between the tumbling correlation times (indicated on each panel) in the dark and lit states is detected, indicative of the same average particle size.

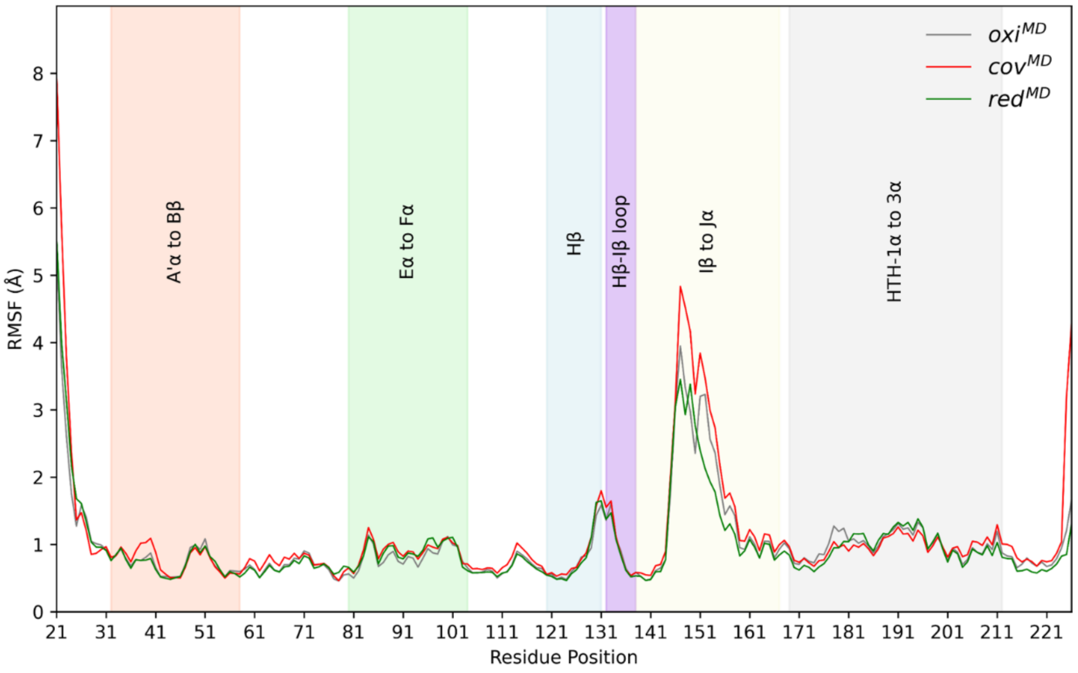

**Fig. S13. Root mean square fluctuations (RMSF) of EL222 by molecular dynamics (MD) simulations.** RMSF values were calculated using Cα atoms of protein residues of $oxi^{MD}$,$cov^{MD}$ and $red^{MD}$ trajectories (3 µs cumulative MD simulations each).

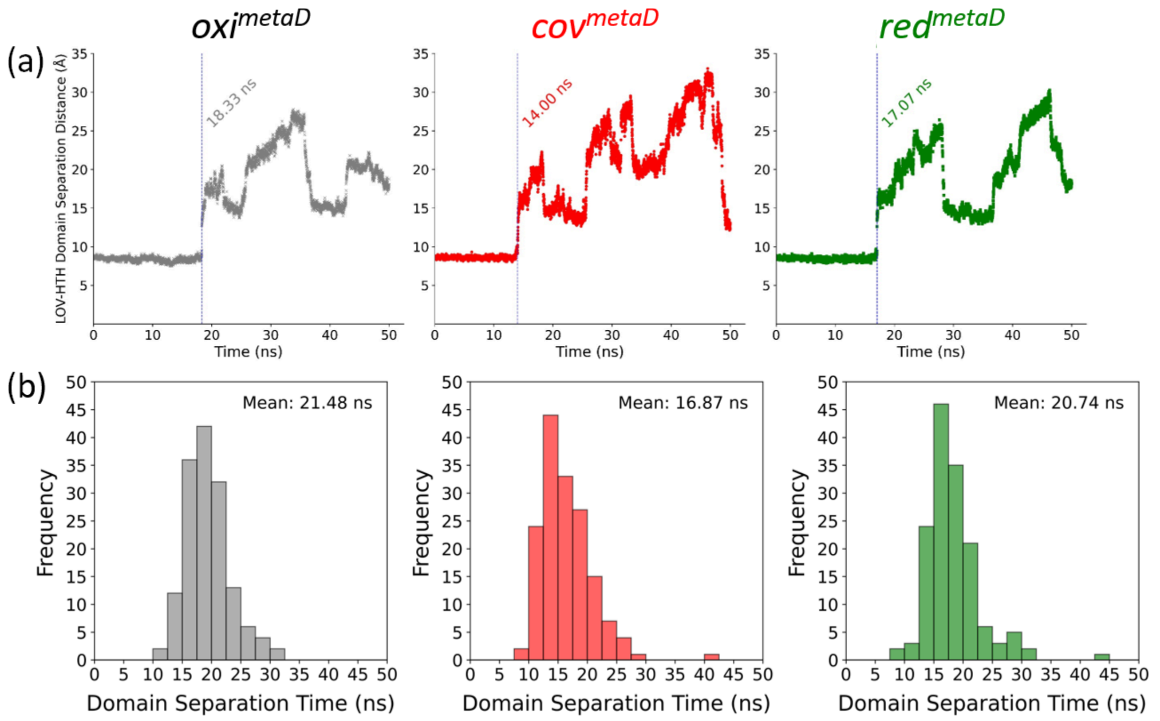

**Fig. S14. Inter-domain opening dynamics of EL222 by metadynamics simulations.** (b) Histogram plots of LOV-HTH domain separation time in ns obtained from metadynamics (*metaD*) simulation (160 replicas * 50 ns length) of *oxi*, *cov*, and *red* starting structures. Mean value for domain separation time is written over the plots. (a) LOV-HTH domain separation distance charges during *metaD* simulation time for $oxi^{metaD}$, $cov^{metaD}$ and $red^{metaD}$ models, sampled from one representative model out of 160 replicas. Vertical dashed lines show the time (in nanoseconds or ns) corresponding to the moment when applied bias effectively caused LOV-HTH domain separation. (b) Histogram plots of LOV-HTH domain separation time in ns obtained from metadynamics (*metaD*) simulation (160 replicas * 50 ns length) of *oxi*, *cov*, and *red* starting structures. Mean value for domain separation time is written over the plots.

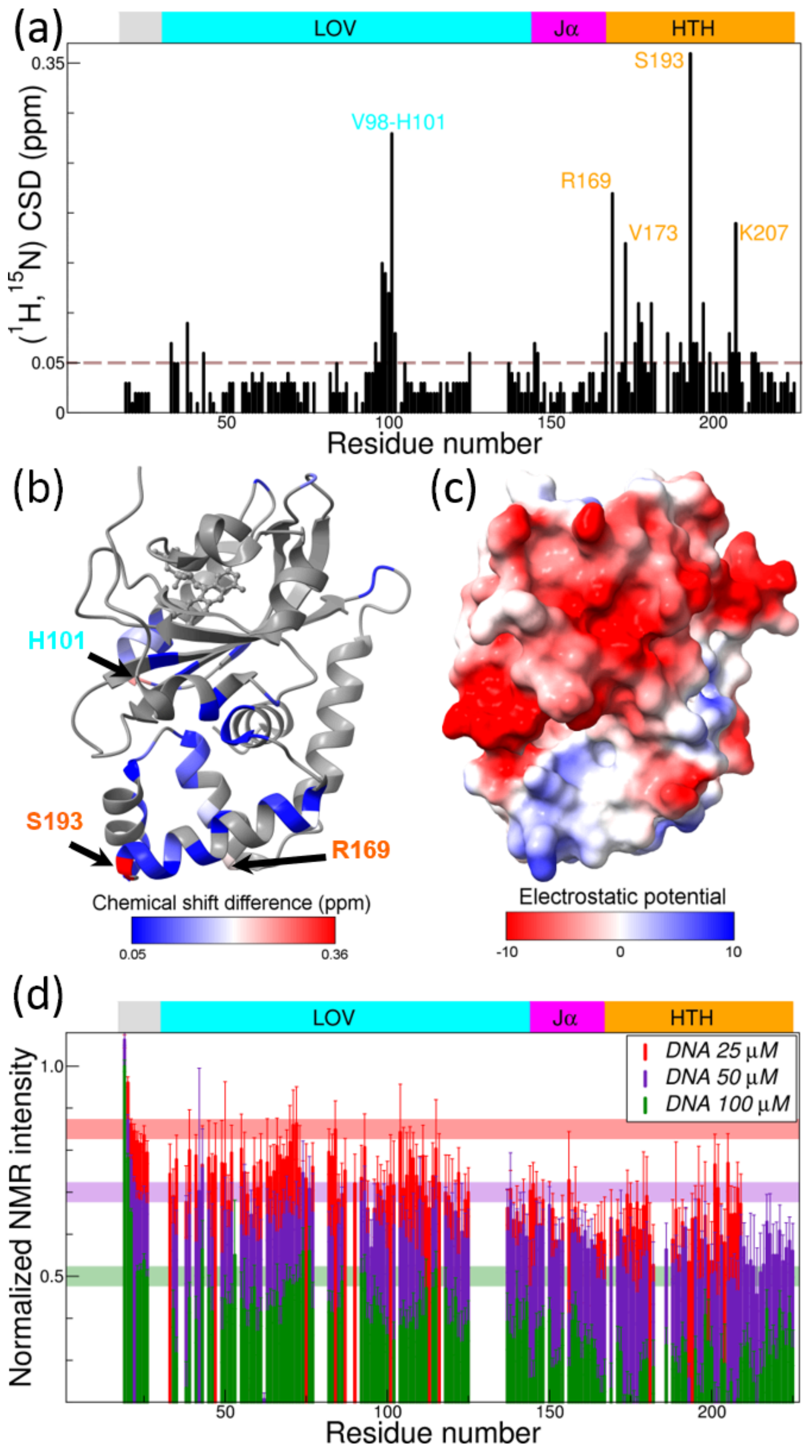

**Fig. S15. Interaction between EL222 and DNA in the dark.** (a) $^{1}H$-$^{15}N$ chemical shift differences (CSD) between EL222-C78A (100 μM ) in presence and absence of 100 μM DNA (dark state). (b) Mapping chemical shift differences from (a) on the dark-state structure of EL222 (PDB ID: 8A5R). For the ease of visualization a threshold of 0.05 ppm was applied. The three residues with the largest chemical shift differences are indicated. (c) Mapping the electrostatic potential on the dark-state structure of EL222 (PDB ID: 8A5R). (d) Drop in EL222-C78A $^{1}H$-$^{15}N$ peak intensity (100 μM) as a function of DNA concentration.

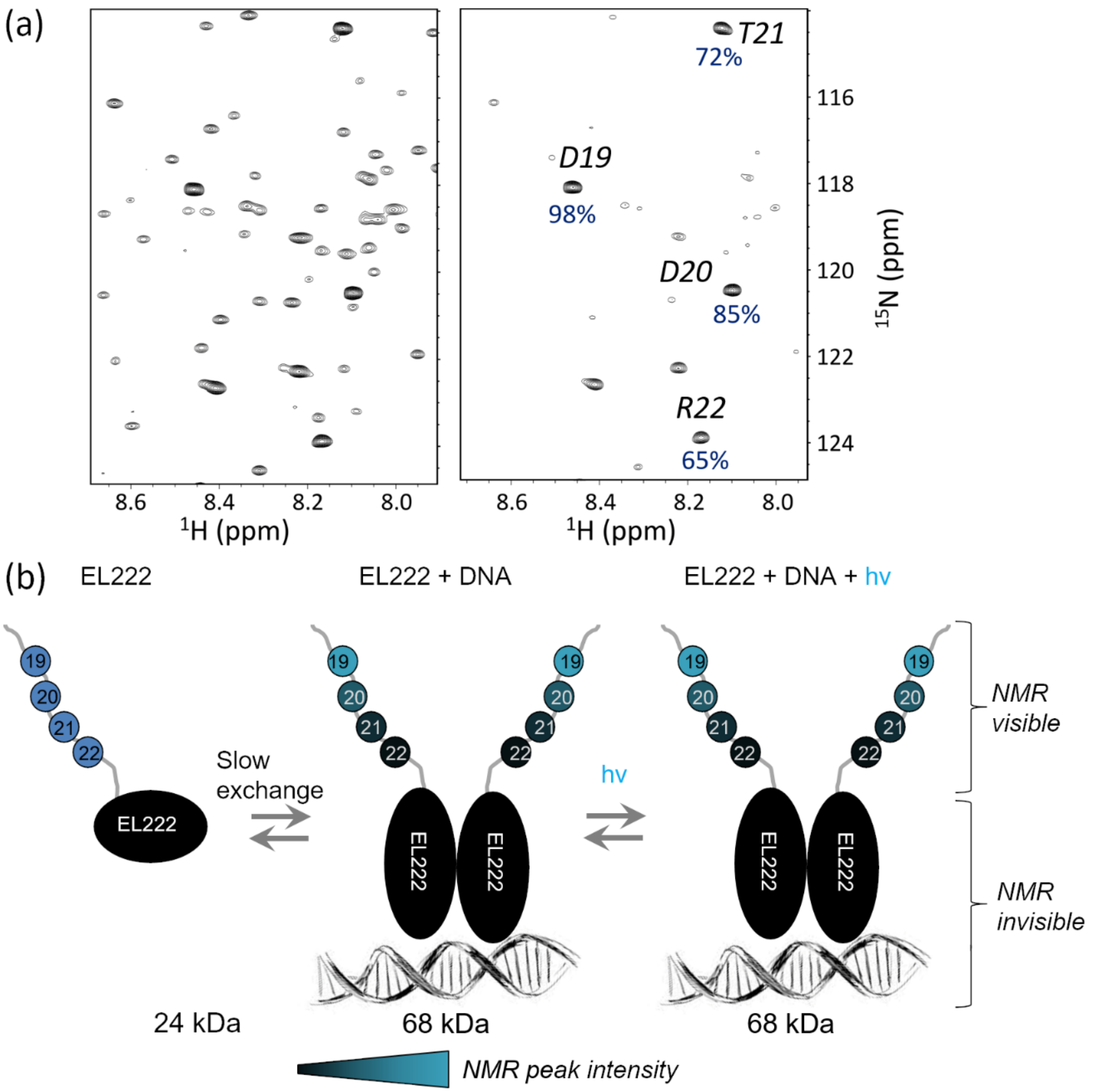

**Fig. S16. Interaction between EL222 and DNA.** (a) $^{1}H$-$^{15}N$ correlation spectra of 135 μM EL222 (left), and in presence of 50 μM DNA recorded in the dark (no illumination) (right). In the presence of DNA, the peak intensity of residues in the LOV and HTH domains decreases to < 65%, rationalized by the formation of EL222:DNA complexes with a 2:1 stoichiometry. The 4 N-terminal residues (Asp19, Asp20, Thr21, and Arg22) show a progressive decrease in NMR peak intensity, indicating that the N-terminus remains highly flexible in the EL222:DNA complex. (b) Cartoon illustrating the interaction between EL222 and DNA. In the dark, EL222 is in a concentration-dependent equilibrium between a free and a DNA-bound form. In the presence of blue light, the equilibrium is shifted towards the EL222-DNA complex.

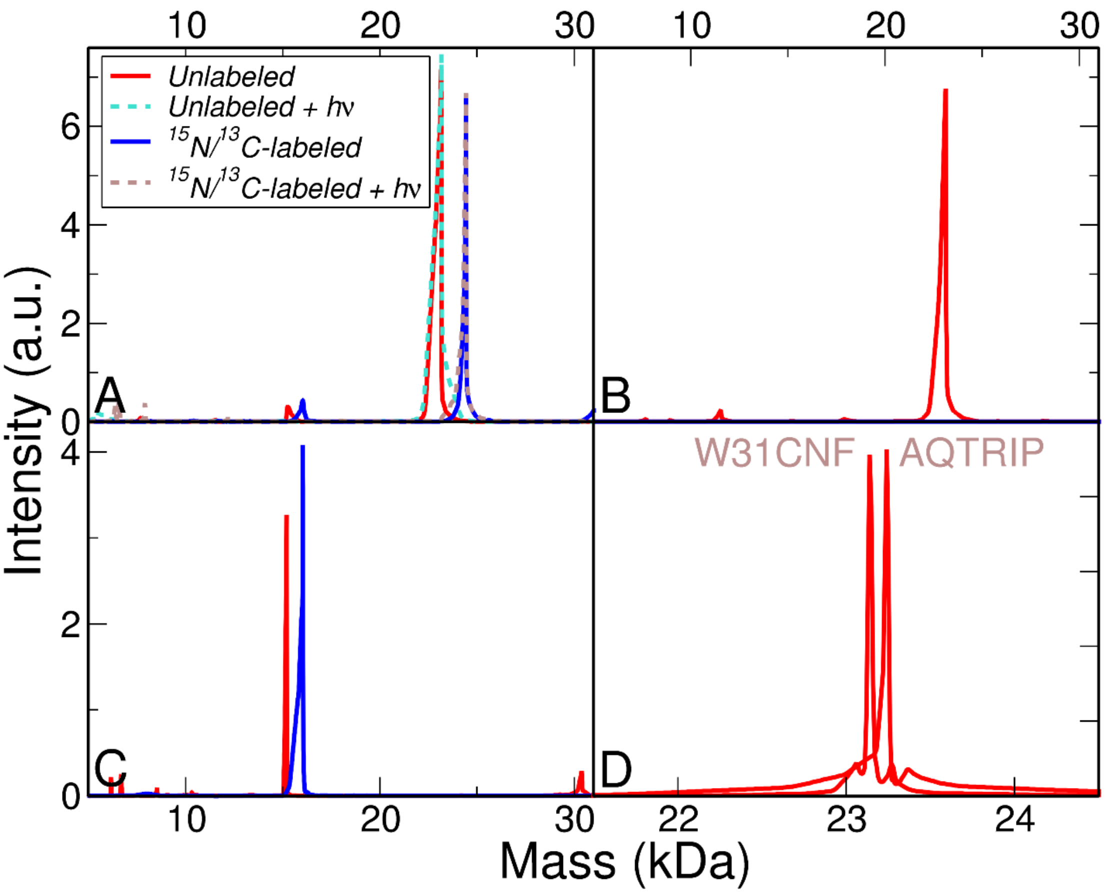

**Fig. S17. Deconvoluted mass spectra of EL222 variants used in this study.** (a) EL222-WT. (b) EL222-C78A. (c) EL222-LOV (d) Others: EL222-W31CNF and EL222-AQTRIP. Unlabeled refers to the normal isotope abundance ($^{14}N/^{12}C$) samples used for IR studies. $^{15}N/^{13}C$-labeled refers to the isotopically-enriched samples used for NMR studies. Samples under continuous exposure to light for several hours are indicated as "+ hv".

**Table S1.** Data collection, processing and model refinement statistics. Values in parentheses are for the highest resolution shell.

| ***Structure*** | *EL222dark* | *EL222light* |
|---|---|---|
| *PDB ID* | 8A5R | 8A5S |
| ***Data collection*** | | |
| *X-ray source* | MetalJet D2 | MetalJet D2 |
| *Wavelength (Å)* | 1.3418 | 1.3418 |
| *Detector* | Photon II | Photon II |
| *Crystal-detector distance (mm)* | 80 | 70 |
| *No. of oscillation images* | 600 | 640 |
| *Exposure time per image (s)* | 30 | 20 |
| *Oscillation width (°)* | 0.3 | 0.25 |
| *Space group* | $P2_12_12_1$ | $P2_12_12_1$ |
| *Unit-cell parameters a, b, c (Å)* | 43.8, 52.0, 81.1 | 43.4, 51.9, 80.4 |
| *Resolution range (Å)* | 30.96–1.85 (1.89–1.85) | 43.57–1.85 (1.89–1.85) |
| *No. of observations* | 72136 (1472) | 83711 (4041) |
| *No. of unique reflections* | 15648 (680) | 16065 (955) |
| *Data completeness (%)* | 95.6 (69.6) | 99.7 (99.9) |
| *Average redundancy* | 4.6 (2.2) | 5.2 (4.2) |
| *Average mosaicity (°)* | 0.18 | 0.18 |
| *Average I/σ(I)* | 13.2 (2.5) | 10.9 (1.9) |
| *Solvent content (%)* | 37.3 | 37.3 |
| $R_{merge}$ | 0.070 (0.437) | 0.116 (0.772) |
| $R_{meas}$ | 0.078 (0.567) | 0.129 (0.881) |
| *CC(1/2)* | 0.998 (0.773) | 0.996 (0.636) |
| *Wilson B factor (Å²)* | 10.8 | 10.9 |
| ***Refinement*** | | |
| $R_{work}$ | 0.160 | 0.161 |
| $R_{free}$ | 0.215 | 0.215 |
| *Average B factor (Å²)* | 18.2 | 17.7 |
| *R.m.s.d. bonds from ideal (Å)* | 0.011 | 0.013 |
| *R.m.s.d. angles from ideal (°)* | 1.727 | 1.688 |
| *Ramachandran favoured (%)* | 99 | 96 |
| *Ramachandran outliers (%)* | 0 | 0 |
| ***Model*** | | |
| *No. of modeled amino acids* | 206 | 206 |
| *No. of modeled alt. conformations* | 17 | 26 |
| *No. of modeled water molecules* | 251 | 223 |
| *Ligands* | 1 x FMN, 1 x MES, 2 x $Cl^-$, 2 x $Mg^{2+}$ | 1 x FMN, 1 x MES, 2 x $Cl^-$, 3 x $Mg^{2+}$ |

**Table S2.** Comparison between the three crystal structures of EL222. Structural superposition of light activated transcription factor EL222 from *Erythrobacter litoralis* (EL222 Dark, EL222 Lit and 3P7N) were performed with the least squares superposition function in O (6) using the default distance cut-off limit of 3.8 Å.

| *Model I, residues* | *Model II* | *No. of aligned amino acids* | *r.m.s.d.* |
|---|---|---|---|
| *EL222 Dark, all CA* | EL222 Lit | 21-226/ 206 aa | 0.217 |
| *EL222 Dark, LOV domain* | EL222 Lit | 38-149/ 112 aa | 0.129 |
| *EL222 Dark, Jα-HTH domain* | EL222 Lit | 167-223/ 57 aa | 0.179 |
| *EL222 Dark, all CA* | 3P7N A | 188 aa | 1.254 |
| *EL222 Dark, all CA* | 3P7N B | 188 aa | 1.259 |
| *3P7N A* | 3P7N B | 191 aa | 0.621 |

**Table S3.** Kinetics of lit1-to-lit2 photoconversion expressed as lit1 lifetimes in minutes.

| | Technique | |
|---|---|---|
| ***EL222 variant*** | ***NMR***[a] | ***IR***[c] |
| *EL222-WT* | 200[b]<br>2200[c] | 160[e] (Amide I' + C=O FMN)<br>320[f] (S-H) |
| *EL222-LOV* | 20[d] | N.D. |
| *EL222-W31CNF* | N.D | 130[f] (C≡N) |

[a] $H_2O$:$D_2O$ (95:5)
[b] 70 mW
[c] 8 mW
[d] 50 mW
[e] 100 %$D_2O$
[f] 100% $H_2O$

**Table S4.** Key restrains used to generate the molecular models for computer simulations shown in **Fig. 4a.**

| ***MODEL*** | ***FMN-C78 adduct?*** | ***Q141 rotation?*** | ***FMN protonation*** |
|---|---|---|---|
| *Oxi* | NO | NO | $N_5 + N_1$ |
| *Cov* | YES | YES | $N_5H + N_1$ |
| *Red* | NO | NO | $N_5H + N_1^-$ |

**Table S5.** Polymer sequences used in this study.

| *Name* | *Sequence* |
|---|---|
| *EL222-WT* | GADDTRVEVQPPAQWVLDLIEASPIASVVSDPRLADNPLIAINQAFTDLTGYSEEECVG RNCRFLAGSGTEPWLTDKIRQGVREHKPVLVEILNYKKDGTPFRNAVLVAPIYDDDDELL YFLGSQVEVDDDQPNMGMARRERAAEMLKTLSPRQLEVTTLVASGLRNKEVAARLGLSEK TVKMHRGLVMEKLNLKTSADLVRIAVEAGIA |
| *EL222-C78A* | GADDTRVEVQPPAQWVLDLIEASPIASVVSDPRLADNPLIAINQAFTDLTGYSEEECVG RNARFLAGSGTEPWLTDKIRQGVREHKPVLVEILNYKKDGTPFRNAVLVAPIYDDDDELL YFLGSQVEVDDDQPNMGMARRERAAEMLKTLSPRQLEVTTLVASGLRNKEVAARLGLSEK TVKMHRGLVMEKLNLKTSADLVRIAVEAGIA |
| *EL222-LOV* | GEFGADDTRVEVQPPAQWVLDLIEASPIASVVSDPRLADNPLIAINQAFTDLTGYSEEEC VGRNCRFLAGSGTEPWLTDKIRQGVREHKPVLVEILNYKKDGTPFRNAVLVAPIYDDDDE LLYFLGSQVEVDDDQPN |
| *EL222-W31CNF* | GADDTRVEVQPPAQ*VLDLIEASPIASVVSDPRLADNPLIAINQAFTDLTGYSEEECVG RNCRFLAGSGTEPWLTDKIRQGVREHKPVLVEILNYKKDGTPFRNAVLVAPIYDDDDELL YFLGSQVEVDDDQPNMGMARRERAAEMLKTLSPRQLEVTTLVASGLRNKEVAARLGLSEK TVKMHRGLVMEKLNLKTSADLVRIAVEAGIA |
| *EL222-AQTRIP* | GADDTRVEVQPPAQWVLDLIEASPIASIVSDPRLADNPIIAINQAFTDLTGYSEEECVG RNCRFLQGSGTEPWLTDKIRQGVREHKPVLVEILNYKKDGTPFRNAVLIAPIYDDDDELL YFLGSQVEVDDDQPNMGMARRERAAEMLKTLSPRQLEVTTLVASGLRNKEVAARLGLSEK TVKMHRGLVMEKLNLKTSADLVRIAVEAGIA |
| *DNA* | 5′-TTATAGGTAGCCTTTAGTCCATGCTGATTCGTT-3′ (forward strand)<br>5′-AACGAATCAGCATGGACTAAAGGCTACCTATAA-3′ (reverse strand) |

*= 4-cyano-L-phenylalanine (CNF)

**Movie S1 (separate file).** <LIT2recovery.MOV>. EL222 recovery to the dark state after prolonged illumination in a NMR (Shigemi) tube.
